# microRNAs (miR 9, 124, 155 and 224) transdifferentiate macrophages to neurons

**DOI:** 10.1101/2020.07.19.210633

**Authors:** Naveen Challagundla, Reena Agrawal-Rajput

## Abstract

Development is an irreversible process of differentiating the undifferentiated cells to functional cells. Brain development involves generation of cells with varied phenotype and functions, which is limited during adulthood, stress, damage/degeneration. Cellular reprogramming makes differentiation reversible process with reprogramming somatic/stem cells to alternative fate with/without stem cells. Exogenously expressed transcription factors or small molecule inhibitors have driven reprogramming of stem/somatic cells to neurons providing alternative approach for pre-clinical/clinical testing and therapeutics. Here in, we report a novel approach of microRNA (miR)-induced trans-differentiation of macrophages (CD11b high) to induced neuronal cells (iNCs) (neuronal markers high-Nestin, Nurr1, Map2, NSE, Tubb3 and Mash1) without exogenous use of transcription factors. miR 9, 124, 155 and 224 successfully transdifferentiated macrophages to neurons with transient stem cell-like phenotype. We report trans differentiation efficacy 18% and 21% with miR 124 and miR 155. *in silico* (String 10.0, miR gator, mESAdb, TargetScan 7.0) and experimental analysis indicate that the reprogramming involves alteration of pluripotency genes like *Oct4, Sox2, Klf4, Nanog* and pluripotency miR, *miR 302*. iNCs also shifted to G0 phase indicating manipulation of cell cycle by these miRs. Further, CD133+ intermediate cells obtained during current protocol could be differentiated to iNCs using miRs. The syanpsin^+^ neurons were functionally active and displayed intracellular Ca^+2^ evoke on activation. miRs could also transdifferentiate bone marrow-derived macrophages and peripheral blood mononuclear cells to neuronal cells. The current protocol could be employed for direct *in vivo* reprogramming of macrophages to neurons without teratoma formation for transplantation and clinical studies.

**Highlights:**

- miR 9, miR 124 and miR155 could reprogramme macrophages to mature neurons.
- miR-induced neuronal reprogramming involves stem cell like intermediate phenotype.

**Graphical Abstract:** Macrophages transfected with miR 9, 124, 155 and 224 alter pluripotency genes and neuronal differentiation genes via various mechanisms as elucidated. NIM components may also manipulate driving neuronal differentiation gene expression inducing formation of neuronal cells.

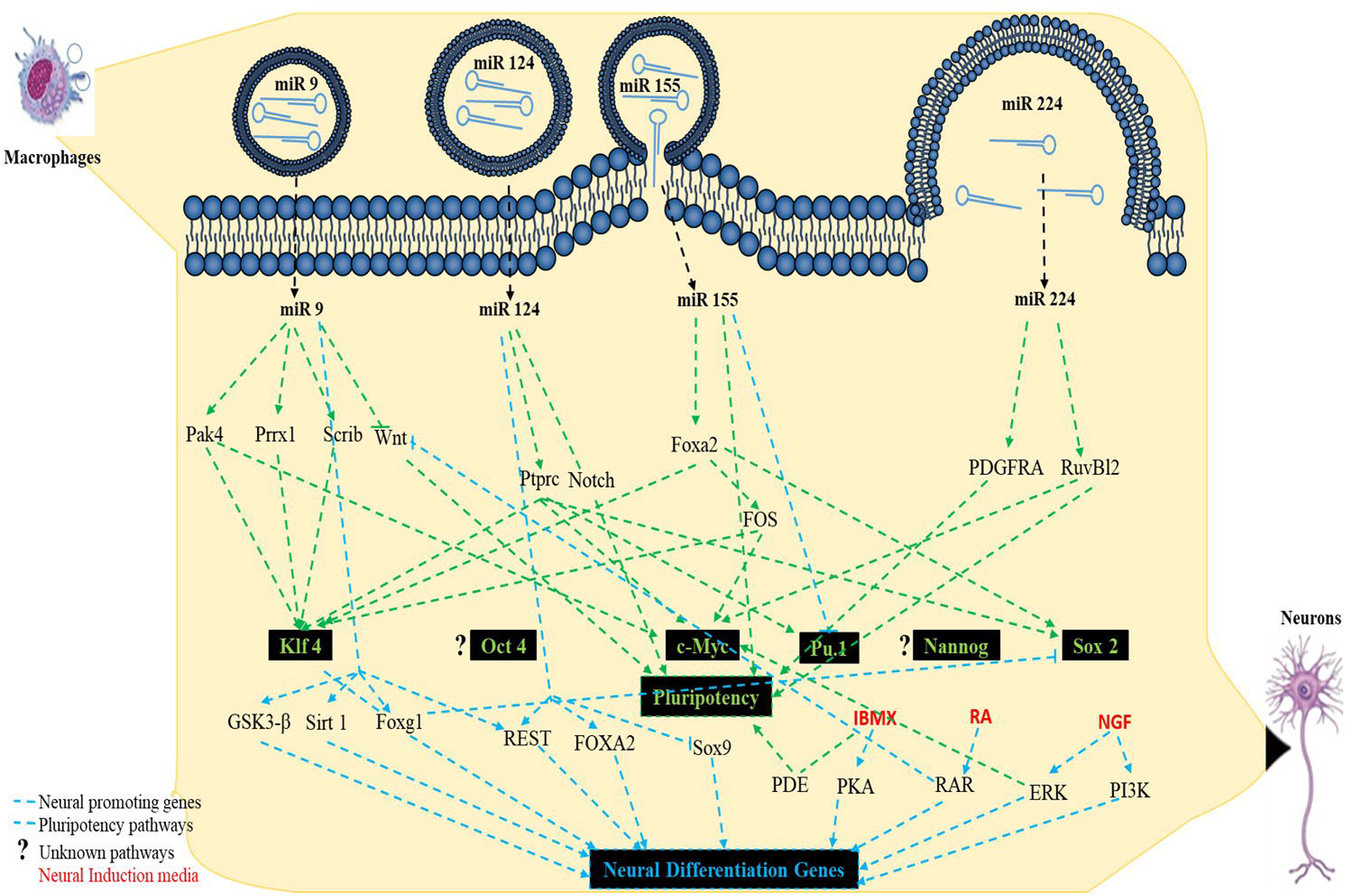

## Introduction

Reprogramming somatic cells into alternative cell type raises hope for obtaining tissues that are difficult to obtain like neuronal tissues. Even after a decade of research on cellular reprogramming significant limitations exist with current protocols thus novel approaches of reprogramming with high efficiency and efficacy are desired. Recent approaches have established a paradigm for forced expression of lineage specific factors to direct reprogramming of somatic cells into desired cell fate with or without induced-pluripotent stem cells (iPSCs). However, these protocols have limitations and remains inefficient owing to the requirement of expressing multiple transcription factors and also due to difficulty in obtaining the source cells. Direct reprogramming of somatic cells is a better strategy compared to reprogramming of stem cells. Reprogramming of fibroblasts, osteoclasts and adipocytes to neurons has been achieved. The study was conceived to transdifferentiate macrophages to neurons considering the following facts. Firstly, macrophages and neurons share phenotypic and functional plasticity and homology (Eyo and Wu, 2013). Secondly, like neurons, macrophages following excitation, transduce signals and display interdependence in some tissues. For example, neurons contribute to macrophage maintenance through release of CSF1, while the BMP2 released by macrophages help in neuronal differentiation (Verheijden et al., 2015). Thirdly, the ease of obtaining macrophages from blood and other tissues makes it an attractive cell target. Lastly and importantly, peripheral macrophages can cross the compromised blood brain barrier (BBB) to reach the site of injury and the resident macrophages (microglia) can easily be reprogrammed to neurons in the brain (Danielyan et al., 2014). Reprogramming macrophages to neurons have thus an edge over other cell types and can easily be translated to pre-clinical and clinical platforms for testing of personalised medicines or transplantation.

MicroRNAs (miRs) are evolutionarily conserved, non-coding small RNAs that regulate the stability and translation of mRNA or subsequent degradation of an active mRNA. miRs have been attributed to play a significant role in regulating gene expression of crucial pathways including but not limited to proliferation, differentiation, apoptosis and metabolism. miRs are the products of long RNA transcripts encoded by miR genes that form hairpin precursors and have been predicted to control and target at least 30% of the entire genome (Faraoni et al., 2009). miR expression is stage- and tissue-specific. miRs regulate many activates in a cell including cell fate determination, stem cell renewal, neurogenesis, plasticity and homeostasis. miR-induced reprogramming has been emerging recently owing to less side effects, wider targets and non-integrative nucleic material. Some reports indicate miR - mediated reprogramming of somatic cells to other cell types. miRs that have been elucidated are preferentially expressed in embryonic stem cells and help maintaining ES phenotype (Judson et al., 2009). Recently miR 307 has been shown to have a role in reprogramming by upregulating OSK factors (Anokye-Danso et al., 2011) while let-7b has been in lime light for reprogramming via regulation of OCT-4, Nanog and SOX2. miRs, along with reprogramming factors enhances the efficacy of reprogramming. Fibroblasts have been reprogrammed to neurons using Bcl2-miR 9-miR 124 overexpressing plasmid (Victor et al., 2014). Most of the miRs reported, induce pluripotency genes and utilize at least one pluripotent transcription factor for efficient iPSC generation (Judson et al., 2009). The present study for the first time, report that miR mimics, without the use of cell cycle or pluripotency transcription factors, can induce cellular reprogramming with better efficacy than the existing protocols.

## RESULTS

miR screening was done for tissue specific miRs using miRGator, EMBL-EMI, mESAdb and miRTargetLink that uses transcriptome analysis (Supplementary fig. S1). Literature suggested that miR 9, miR 124 and miR 155 were increased in different tissues of brain in both human and mice. We also included miR 224 in the present study as it was also earlier reported have a key role in differentiation of neurons, adipocytes, arteries and skin.

### miR transfection and its validation

Manipulation of miR expression in cells is necessary to understand how miRs regulate cell fate. Several methods have been proposed to overexpress miRs. However, overexpression by vectors with small hairpin RNA cassettes or endogenous pre-miRNA may display off-target effect. miR mimics are substantially smaller, easier to transfect and target mRNA more specifically and are thus used in the study. This method is useful for performing functional screening of known miRs. miRNA 9, 124, 155 and 224 are highly conserved across the species and demonstrate brain specific functions. To determine whether these miRs could reprogram somatic cells, we transfected mouse macrophage, Raw 264.7 (macrophage cell line), bone marrow derived macrophages or PBMCs with miR mimics aiming to transdifferentiate them to neurons. Macrophages were transfected with mentioned miRs for 24 hrs and transfection was confirmed by checking the abundance of these miRs or by analysing the fold change in their target genes by real time PCR and sq-RTPCR. The transfected cells showed a maximum of 18 fold and a minimum of 6-fold increase in availability of transfected miR (Fig. 1B). To validate the functionality of miR mimics, we assayed the experimentally validated miR target genes *sirt1* (Fig. 1C), *lamc1* (Fig. 1D), *FADD* (Fig. 1E) and *Ptx3* (Fig. 1F) of miR 9, miR 124, miR 155 and miR 224 respectively. We report a significant change in the expression levels of miRs targets with a minimum of 40% (Fig. 1D) and a maximum of 80% (Fig. 1C and E) suppression in target genes compared to the non-transfected controls as seen in sq-RTPCR analysis (Fig. 1C-F), thus indicating the efficacy of transfection.

**Fig. 1:**
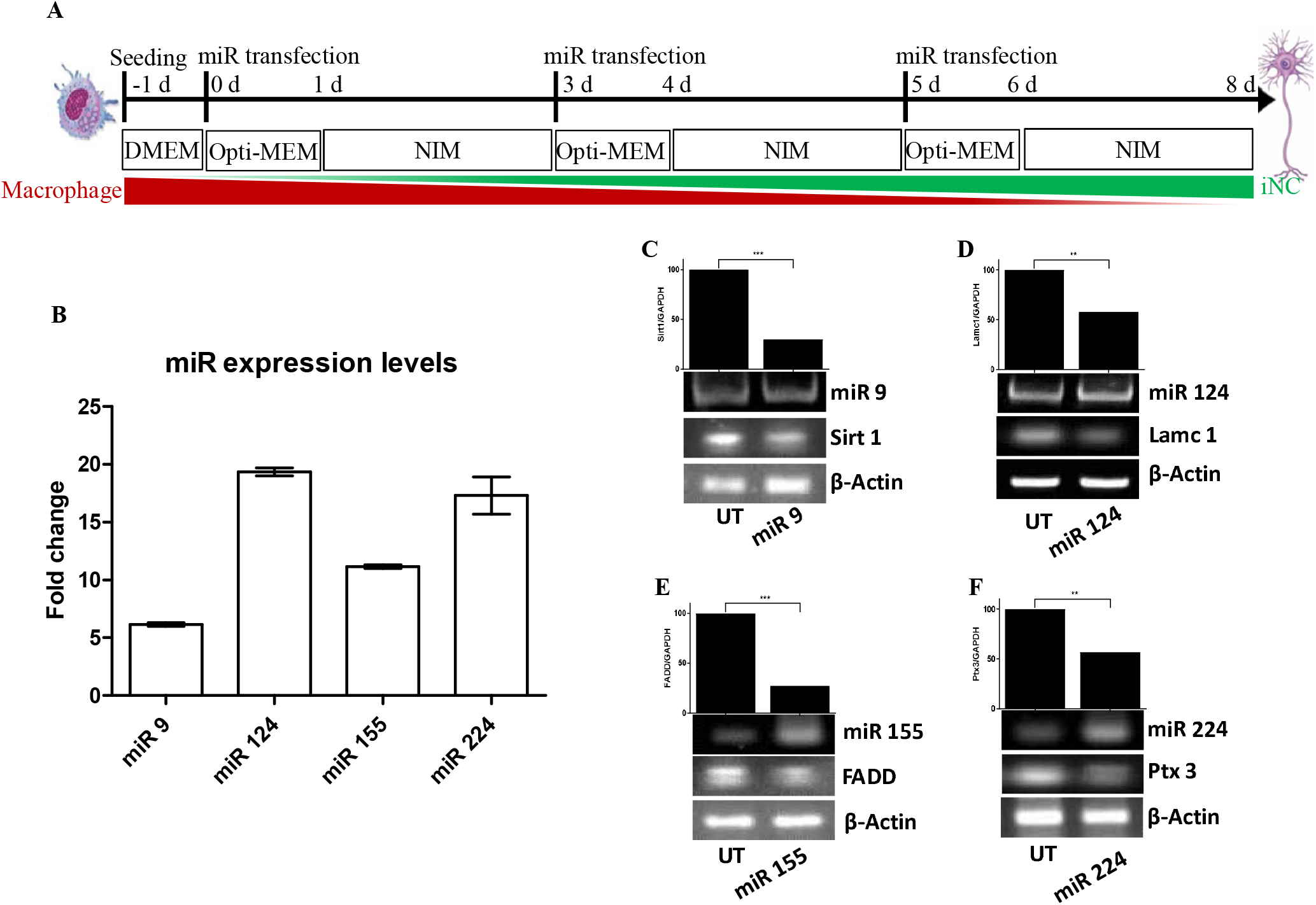
miR transfections and inhibition of target genes. (A) Scheme for miR assisted cellular reprogramming of macrophages. (B) miR transfection was confirmed by Real time-PCR after 24 hrs of transfection. sq-RT PCR analysis of established miR targets (C) *Sirt 1*-miR 9 target; (D) *Lamc1*-miR 124 target; (E) *FADD*-miR 155 target and (F) *Ptx3*-miR 224 in transfected and non-transfected cells. (*** denotes P<0.0001, highly significant; ** denotes P<0.001, Significant)

### miR mimics induce cellular reprogramming and drive macrophages to induced neuronal cells (iNCs)

Macrophages were transfected with miRs and a gradual and significant alteration in the morphology was observed and the reprogrammed cells showed prominent dendrite-like processes that are elongated and branched (Supplementary fig. S2). To confirm the gain or loss of lineage specific markers, the macrophage and neuronal markers were analysed by sq RT-PCR. A substantial decrease in the myeloid marker, *CD11b* (Fig. 2A) and CD11c (Fig. 2B and B’) was observed while the mRNA expression of neuronal marker *nestin* is increased (Fig. 2A). To confirm that the generated neuron-like cells were mature neurons, the generated cells were analysed for neuronal marker expression by RT-PCR. The reprogrammed cells showed an increased expression of *Map2* (Fig. 2C), *Tubb3* (Fig. 2D) and *Mash1* (Fig 2E) in the transfected cells. Map2, a microtubule associated-structural molecule, is specifically expressed in the neurons. Nurr1, a marker for dopaminergic neurons helps in maintenance of their functionality. NSE is an enolase expressed by mature neurons. NeuN is nuclear protein and associated with neuronal differentiation. Synapsin is expressed at functional synaptic vesicles of neurons and play multiple role in transmission, synaptic formation and regulation. Semi-quantitative RTPCR analysis of transfected cells show increase in *NeuN* and *Nerve growth factor receptor (NFGR)* indicating that the cells express neuronal genes (Fig. 2F and F’). Western blot analysis of the neuronal markers, NSE and MAP2 in generated cells confirmed that the reprogrammed cells were indeed neurons. (Fig. 2G and 2G’).

**Fig. 2:**
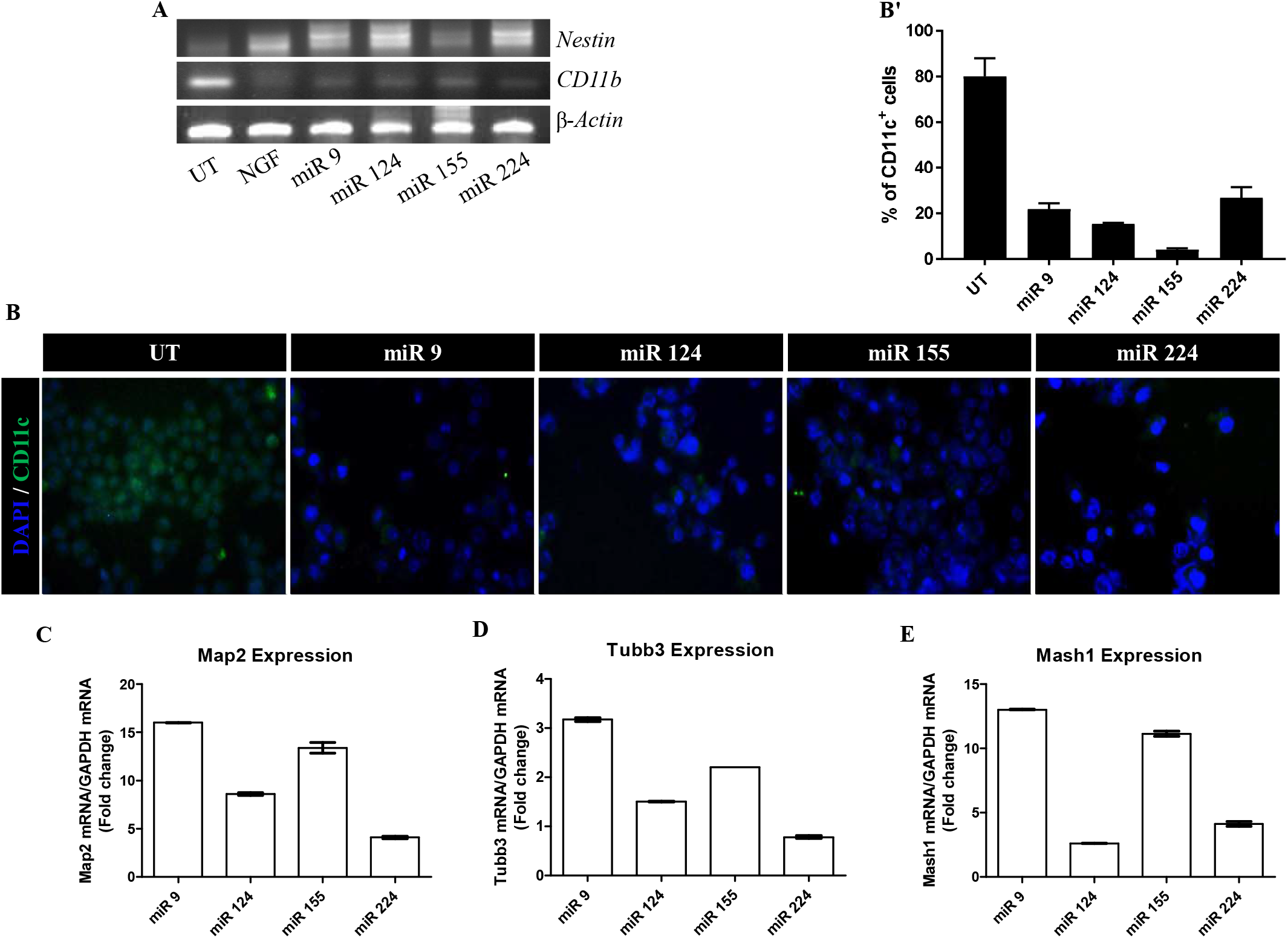
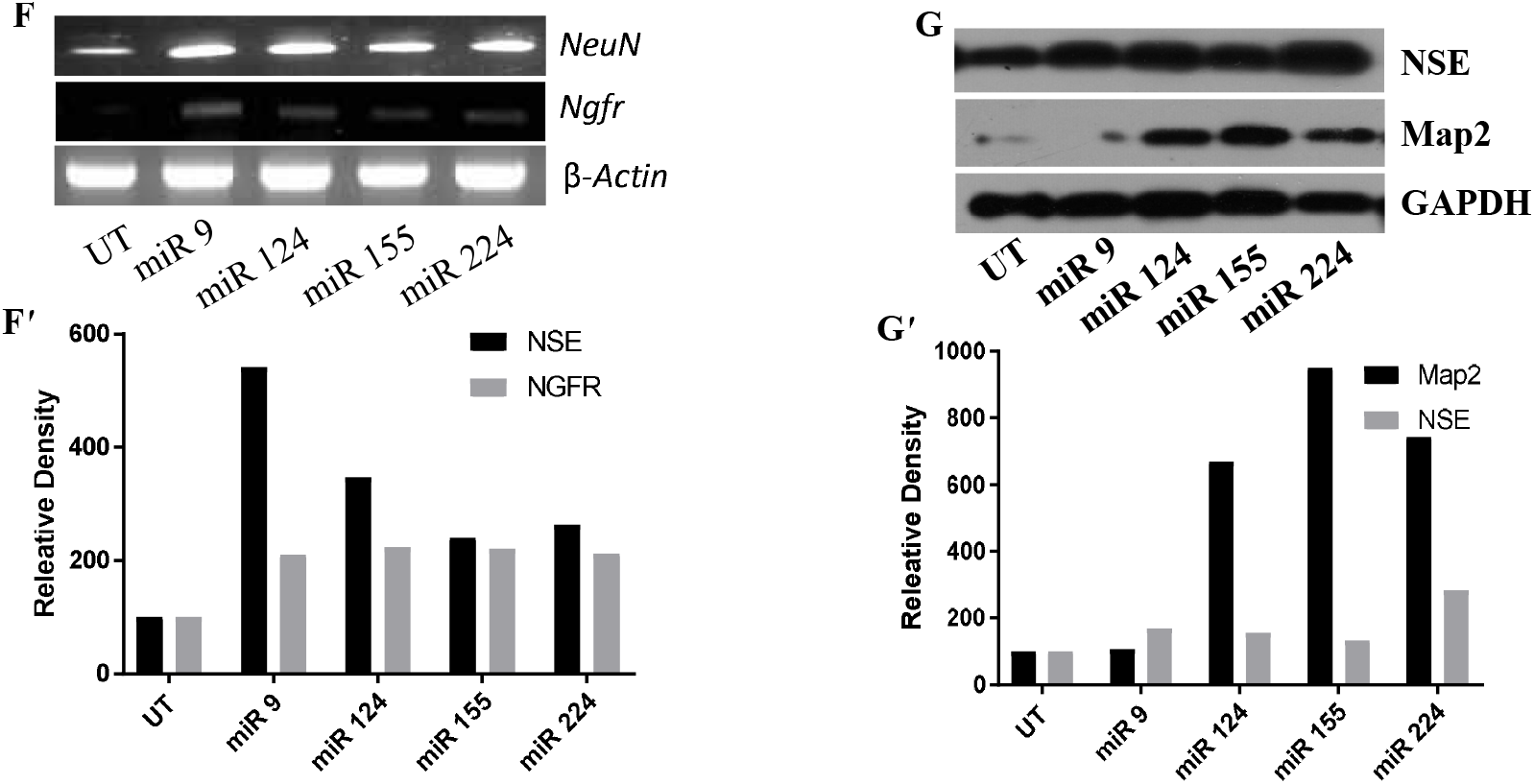
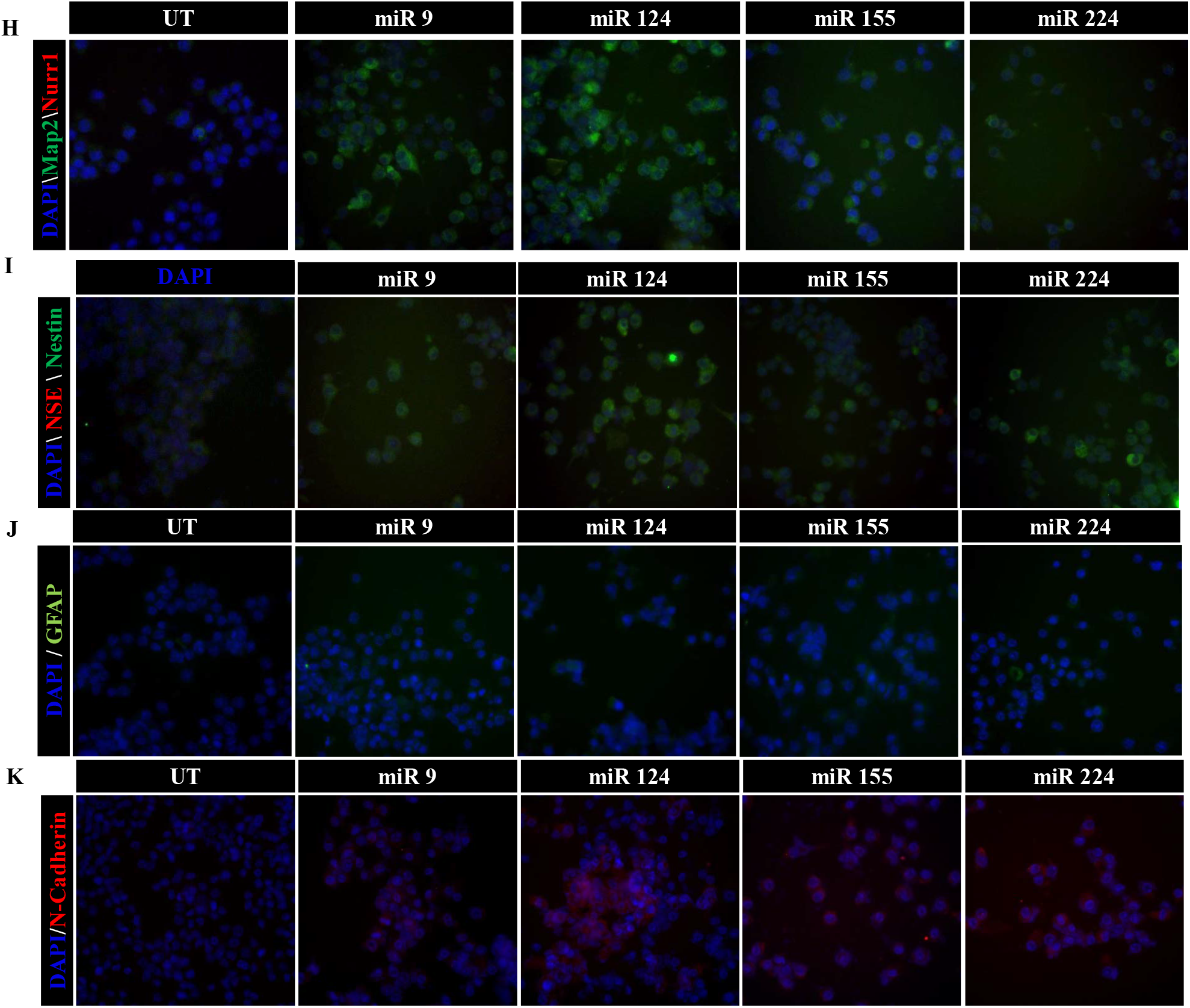
miR mimics transdifferentiate macrophages to induced Neuronal cells (iNCs). (A) Macrophages were transfected with miRs as indicated earlier for three times and gene expression was analysed by sq-RT PCR for *Nestin* and *CD11b*. (B) Immunofluorescence of a myeloid integrin, CD11c of reprogrammed cells. (B’) Cells expressing CD11c were counted in three random fields and plotted as % of CD11c^+^ cells. Real time PCR analysis of Map2 (C), Tubb3 (D) and Mash1 (E) in neuron-like cells. (F and F’) Mature neural marker, *NSE* and *NGFR* were analysed by sq-RT PCR. (G and G’) Increase in neural markers NSE and Map2 was analysed using Western blot in iNCs. iNCs were fluorescently stained with various neuronal markers: (H) Map2 and Nurr1; (I) Nestin and NSE; (J) GFAP and (K) N-Cadherin. (400X magnification). (See supplementary figures S3A, S3B, S3C, S3D aslo) (Relative density was plotted using data from three individual experiments).

Immunofluorescence imaging with neuron specific markers was carried out to see the distribution and expression of markers in individual cells rather than in the pool of cells. Both the early and late neuronal markers were analysed by immunofluorescence. The generated neurons showed an increased expression of early neuronal marker Nurr1 but the fluorescence intensity of Map2 and NSE, mature neuronal markers, were much higher in the transformed cells indicating that the generated cells are mature neurons (Fig. 2H and I; supplementary figure S3A, A’ and A’’). Expression of Nurr1 indicates that the generated iNCs might be dopaminergic neurons. The reprogrammed cells also expressed nestin, a neural stem cell marker (Fig. 2I; supplementary figure S3B, B’ and B’’), which indicates the possibility of reprogramming via neural stem cells. iNCs stained for Glial filamentary acidic protein (GFAP) showed minimal expression (Fig. 2J; supplementary figure S3C). Trans-differentiated neurons expressed validated structural marker N-Cadherin (Fig. 2K; supplementary figure S3D and D’).

### Repeated miR transfection enhance reprogramming efficacy to generate highly pure iNCs

Based on the structural and phenotypic confirmation of transdifferentiated cells we wanted to check the progression of trans-differentiation to get an indication whether it is an immediate and a spontaneous or gradual and systematic process. A kinetic analysis of neuronal marker expression after each transfection was carried out. Neuronal markers were analysed by flow cytometry to assess trans differentiation qualitatively and quantitatively. A gradual increase in the expression of synapsin and Map2 was observed indicating that trans differentiation is systematic process. After three transfections i.e., on day 12, miR 155, miR 124 and miR 224 assisted generation of iNCs were highly pure with more than 98% of reprogrammed cells expressing Map2 and synapsin while with miR 9, it was 93.2% (Fig 3A). To assess the quantitative efficacy of the trans differentiation, we have calculated the percentage generated pure iNCs obtained after each miR transfection with respect to the initial input of macrophages (Fig 3B). The formula used to calculate the efficacy is mentioned in supplementary data. miR transfection could reprogramme macrophages with highest trans differentiation efficacy of 21% as demonstrated with miR 155. (Fig. 3C). miR 124 demonstrated the %age efficacy of 18%; followed by miR 9 which demonstrated the %age efficacy of 16%. The least %age efficacy was demonstrated with miR 224 where only 9% of the input macrophages were transdifferentiated to neurons. With each transfection, the cell number is observed to decrease while the purity is increased. Even though cell number decreases significantly, the purity of the cells follows systematic increasing which infers that the selection process is favourable, particularly for neurons. Despite of repeated transfections the generated and highly pure neurons were viable. The generated neurons were able to maintain the expression of neuronal markers even on 20^th^ day in neurobasal media (Thermo Fisher Scientific) despite withdrawal of miRs and NIM which indicate a stable reprogramming (data not shown).

**Fig. 3:**
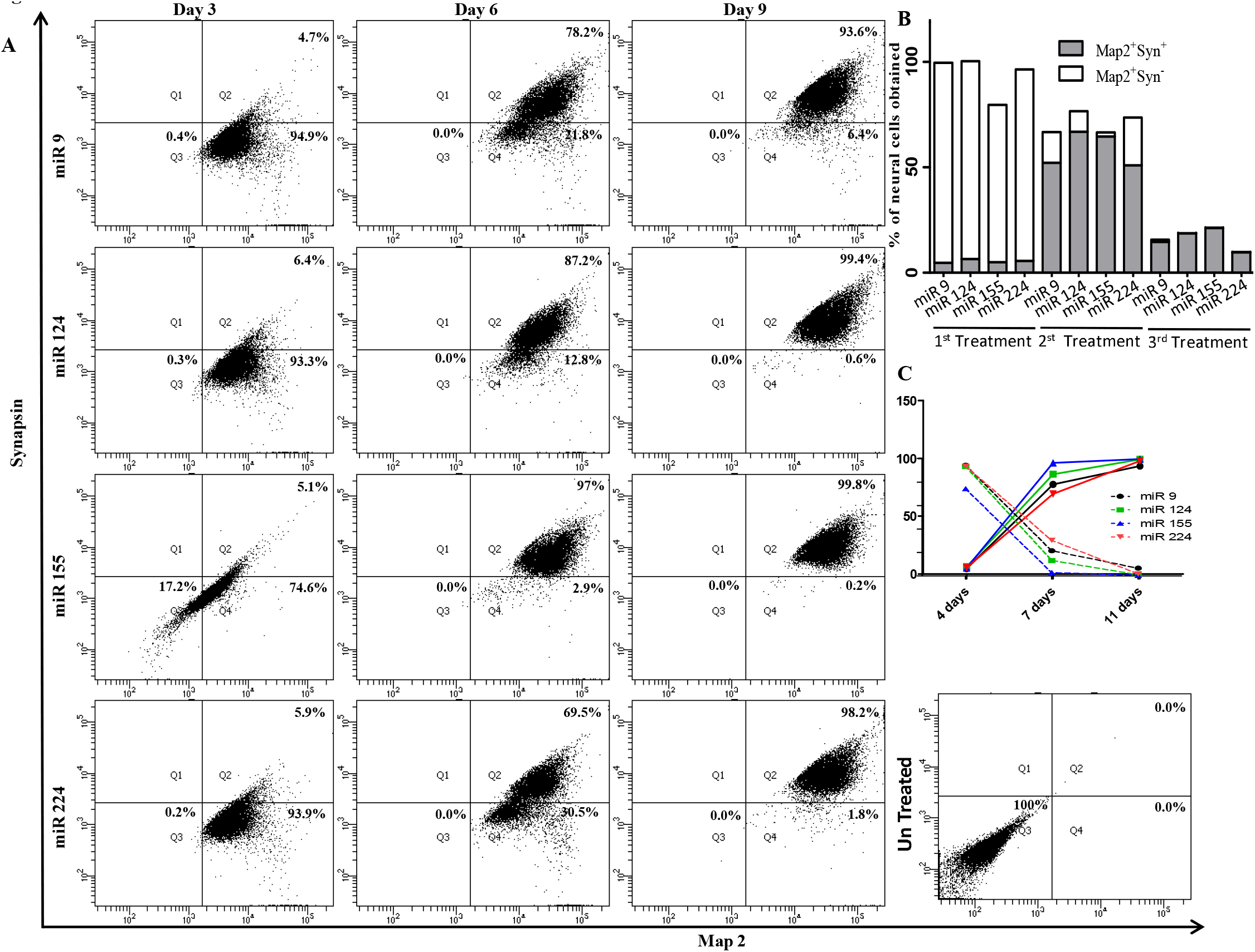
miR transfections transdifferentiate macrophages to pure iNCs. (A) Flow cytometry analysis of 5 x 10^5^ cells were done after every transfection for synapsin and Map2. (B) Efficacy of trans differentiation was plotted calculated from flow cytometry. (C) Trans differentiation efficacy and total no. of iNCs obtained after reprogramming were plotted.

Since, the decrease in cell number might be due to unfavourable conditions for macrophage to survive. The miR used are involved in cell cycle fate manipulation which might inhibit its cell division. We further assessed the cell cycle of generated cells to check the effect of miR on proliferation. The un-transfected cells mostly were observed in the G_1_/S phase indicating active cell division machinery, while, iNCs displayed typical G_0_/G_1_ phase which also explains the loss in cell number after each transfection (supplementary Fig. S4). This further confirms that the cells are differentiated to neurons which generally exist in G_0_ phase. Consistent with our results, it was earlier reported that miR 9 and 124 inhibited Cyclin D and promote cell cycle exit required for neuronal differentiation (Shenoy and Blelloch, 2014).

### miR-mediated reprogramming generated stem cell like-intermediates which are later converted to iNCs

We wanted to elucidate whether the miR mediated reprogramming is direct or via via stem cells, we analysed the relationship between known reprogramming factors and miR targets. Various bioinformatics tools were used for predicting targets and gene ontologies for concentrating their function and pathways. String 10.0 analysis with action details was done using miR predicted targets from the pool of various tools and known reprogramming transcription factors to know the interactions (Fig 4A). Surprisingly, the network of all the miR target genes have shown strong interactions with the validated transcription factors involved in reprogramming (Fig 4B-E).

**Fig. 4:**
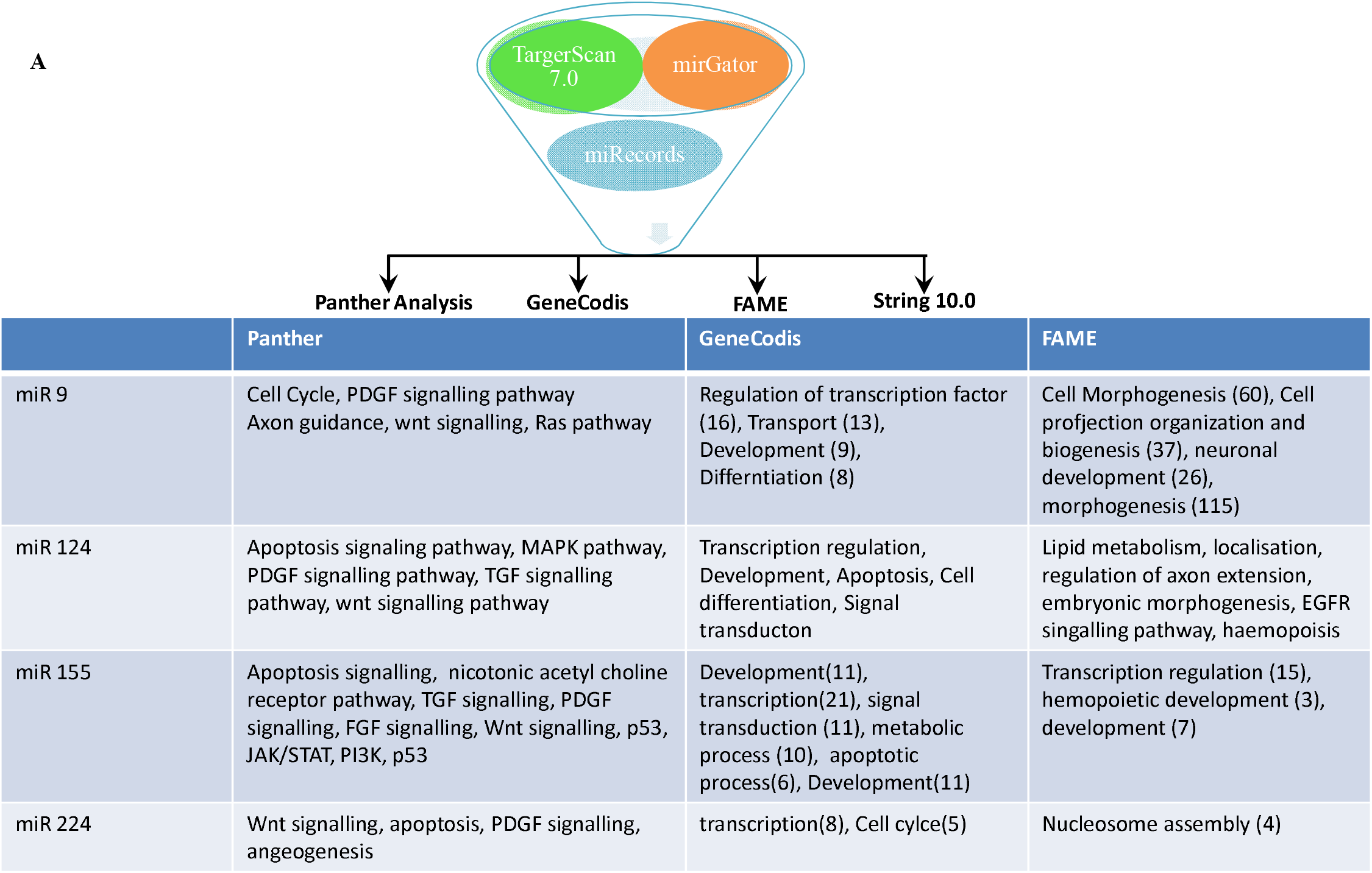
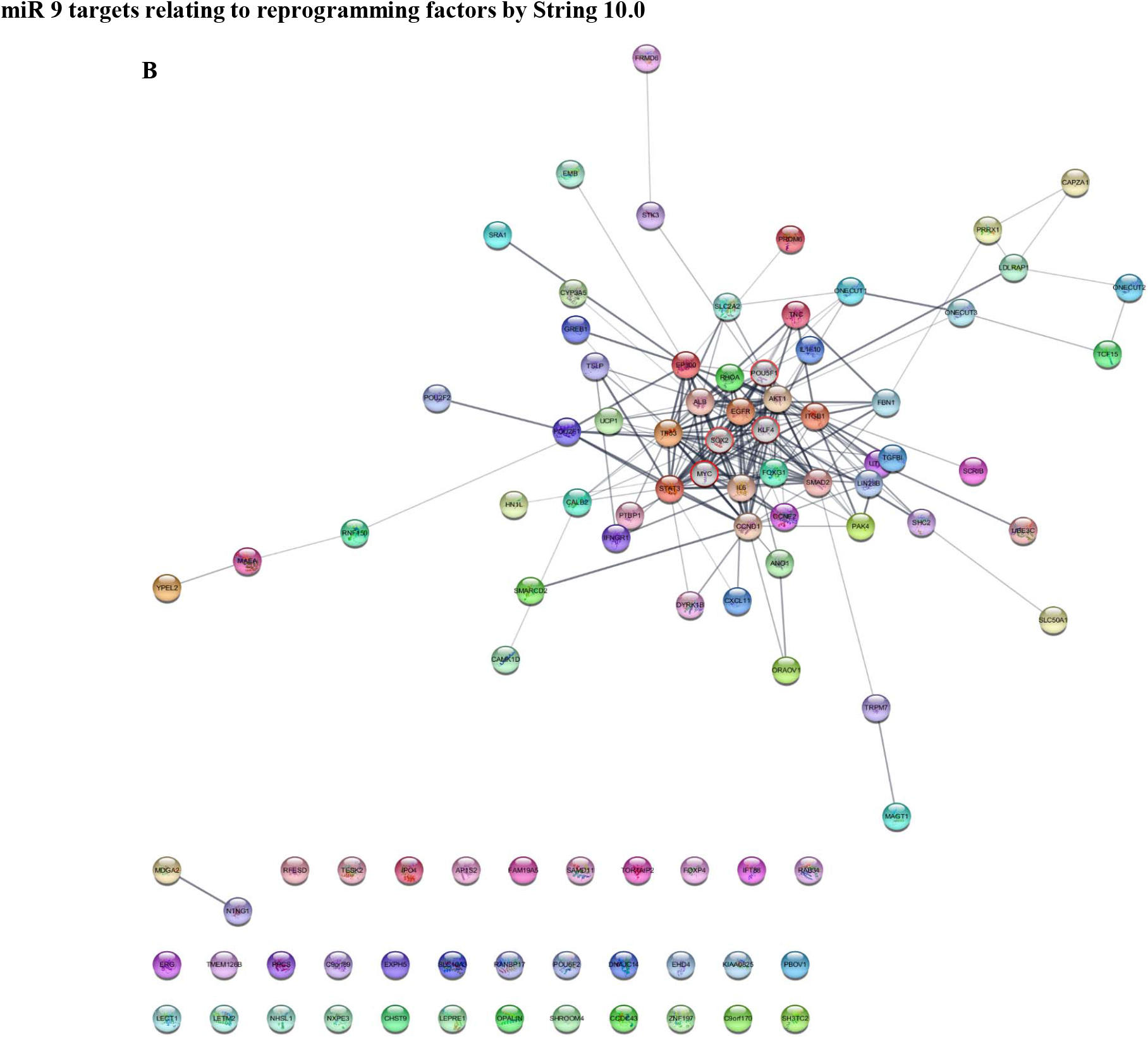
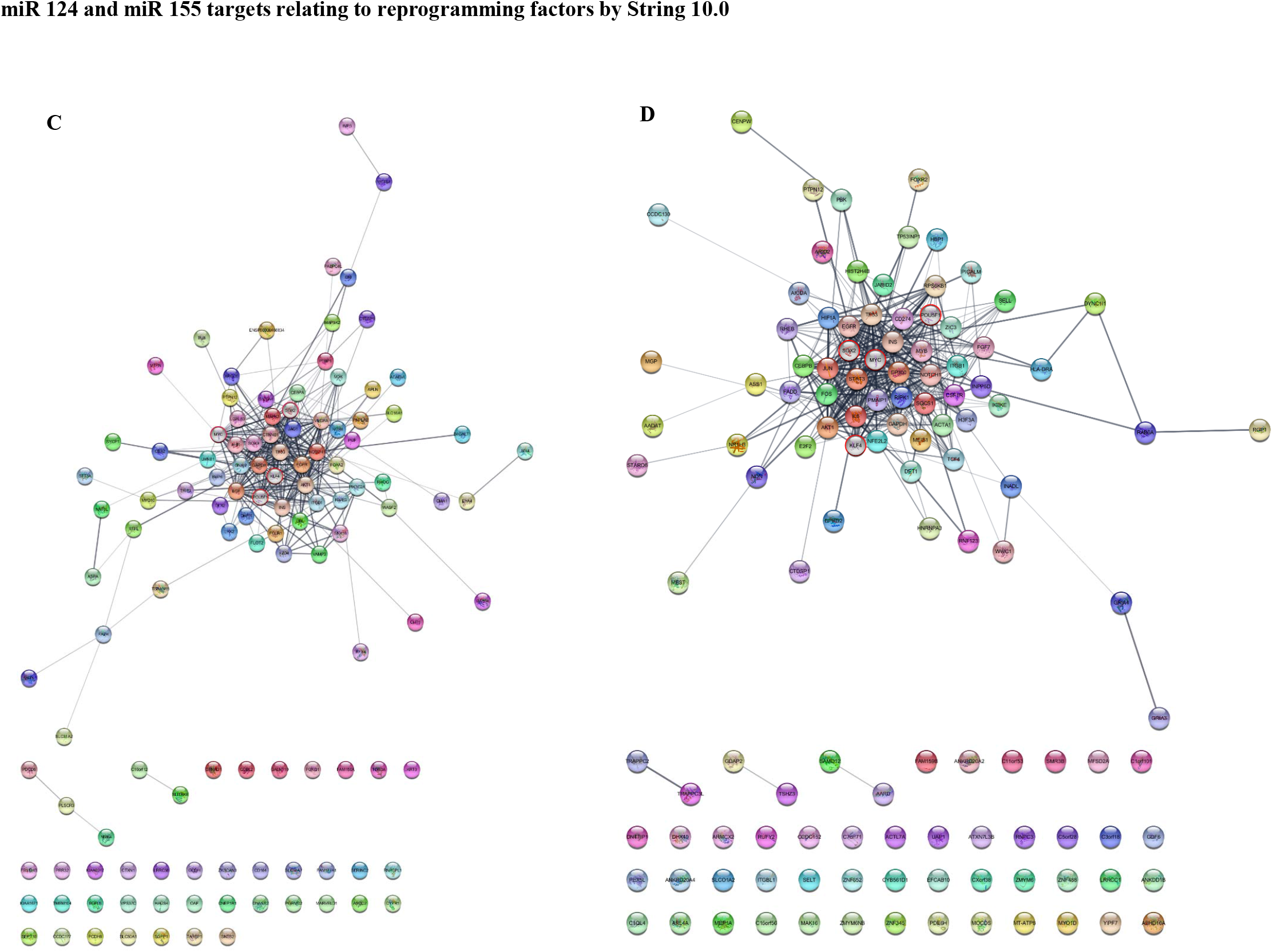
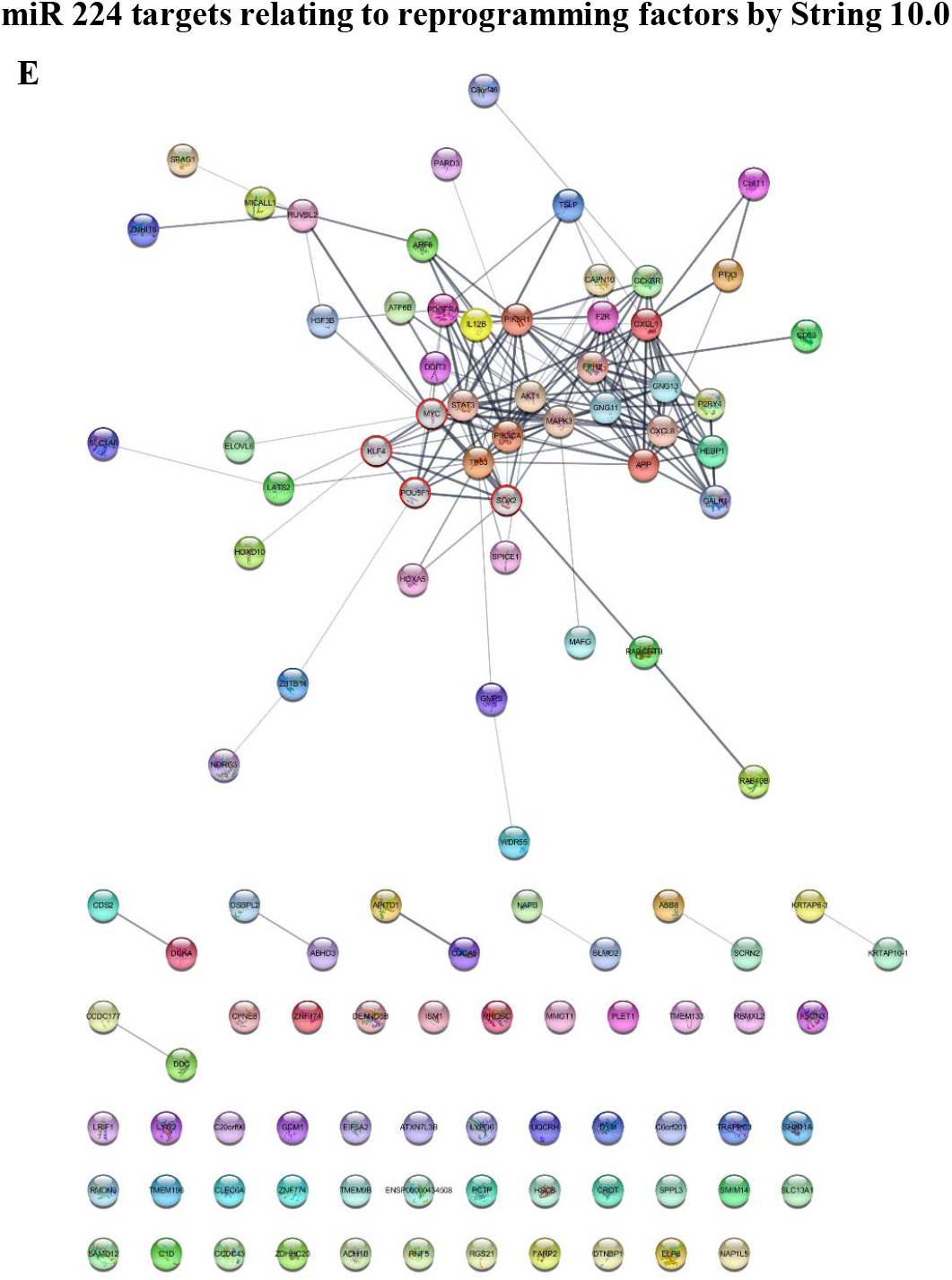
Bio-informatics analysis of miR targets and interaction with pluripotency genes. Gene targets of individual miRs from various databases mentioned were pooled and analysed for their interactions with reprogramming factors and functional ontology. miR targets of miRs were obtained and functional ontology was applied (A). Interaction between (B) miR 9 targets; (C) miR 124 targets; (D) miR 155 targets and (E) miR 224 targets and pluripotency genes was visualised in Cytoscape 3.7.2. OSKM factors were circled in Red and nodes were coloured grey.

Since, the String 10.0 analysis revealed the involvement of pluripotency genes, we wanted to experimentally demonstrate the possibility of involvement of stem-like intermediate cells. The cells displayed increase in the expression of Prominin-1 (CD133) following first transfection. However, after the second and third transfections most of the Prominin^+^ cells express high levels of NSE. Complementing the previous results, miR 124, 155 and 224 show a highly pure iNCs with almost 100% purity. While, miR 9 still had approximately 7% of Prominin^+^ NSE^-^ cells indicating the presence of stem like cells (Fig. 5A). The data thus indicate a strong possibility of cellular reprogramming via an intermediate stem-like cell. The results also co-relate with neural stem cell marker nestin expression (as shown in Figure 2A and I) confirming the involvement of stem like/ neural stem-like cells.

**Fig. 5:**
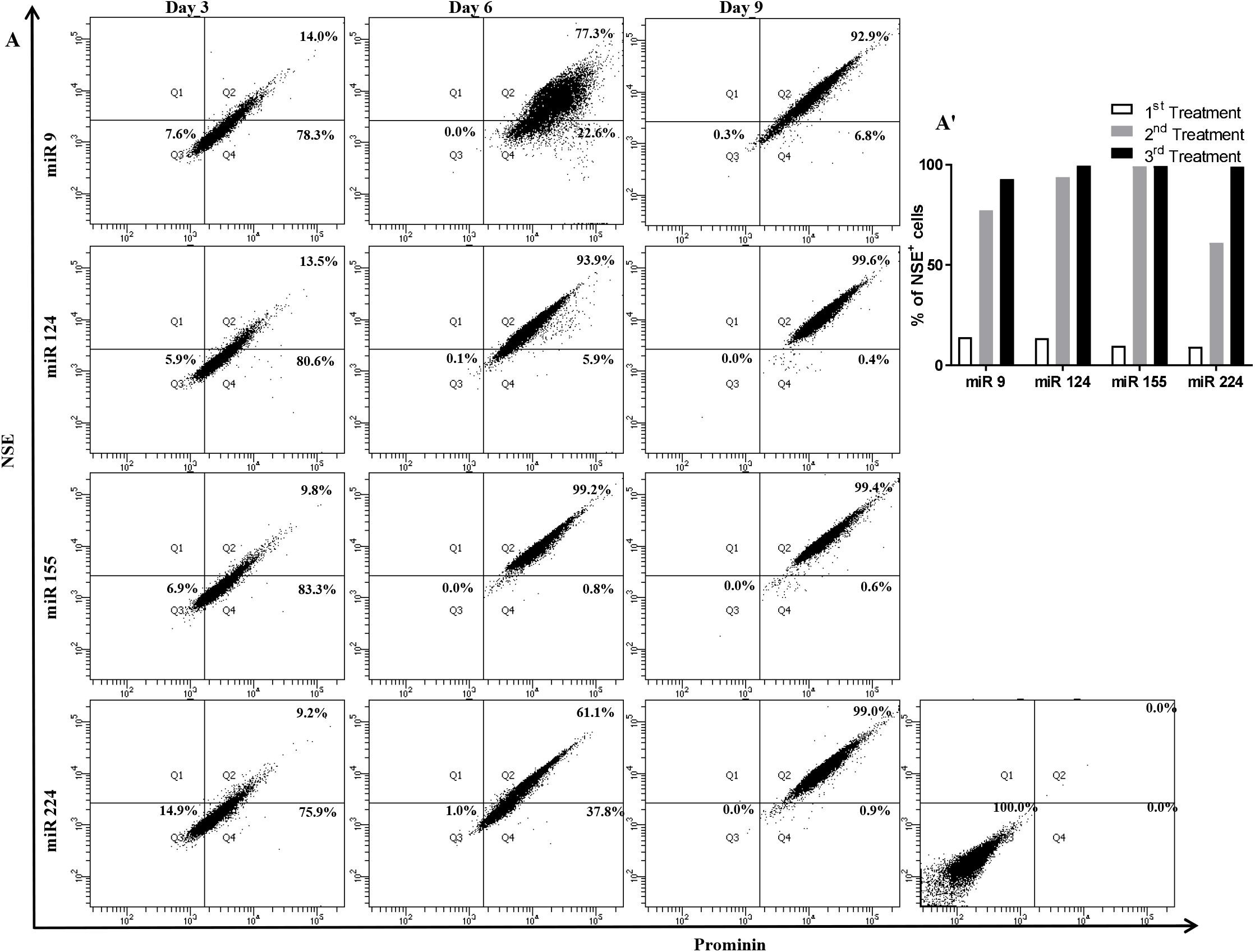
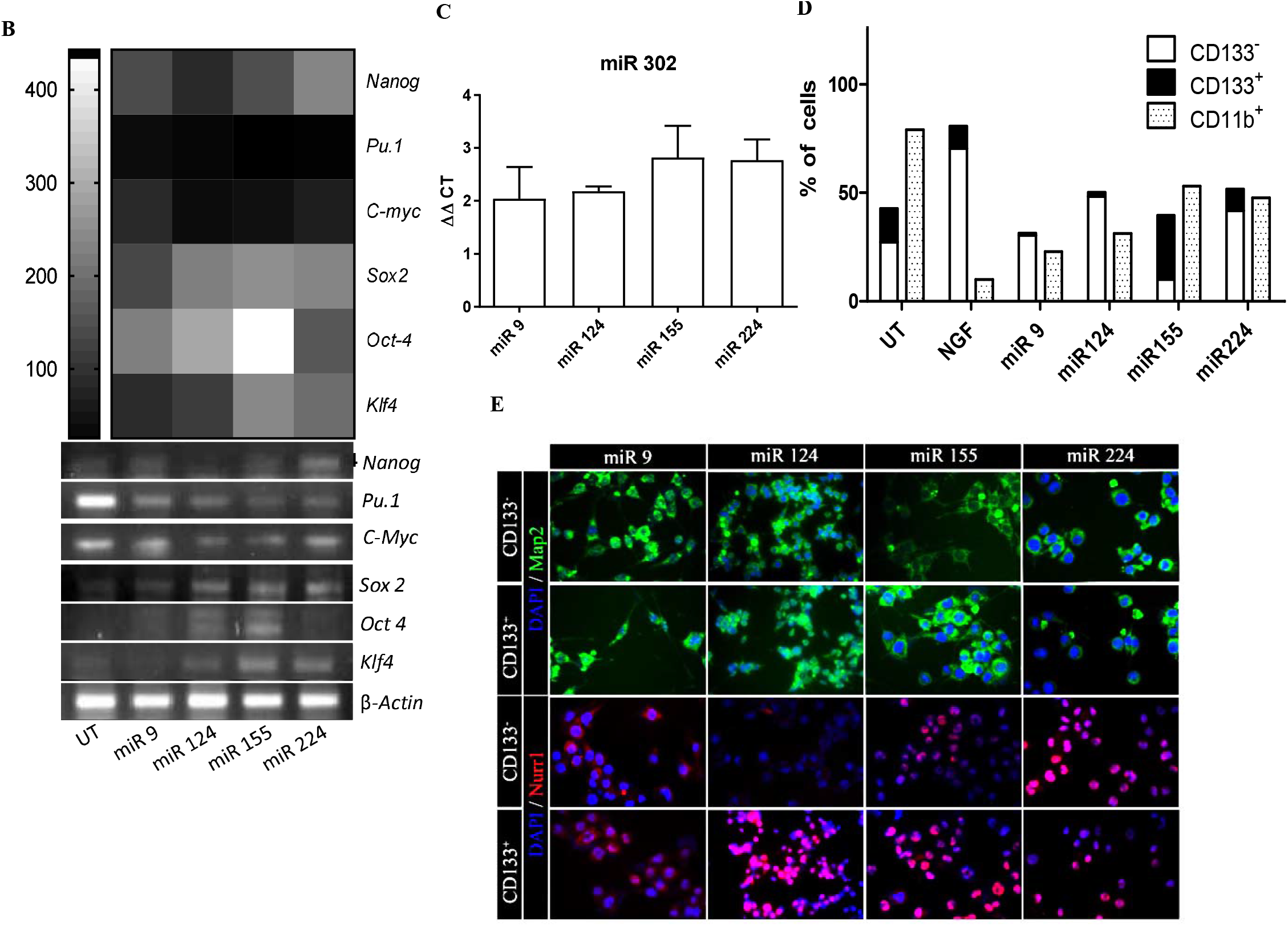
miR-mediated reprogramming involves stem cell-like intermediates. Macrophages were transfected with miRs twice and analysed for Stem cell markers, neuronal marker and pluripotency genes. (A) Flow cytometry analysis of transfected cells were done for NSE and Prominin (A’) NSE expressing cells are represented through histogram (n=3). (B) Relative density of the genes to *β*-actin were calculated and plotted as Heat Map considering *Pu.1* in miR 224 as least and *Oct4* in miR 155 as highest expression (C) miR 302 levels were measured RT-PCR in miR transfected cells. (D) Reprogrammed cells were selected on the basis of expression of CD11b and CD11b^-^ cells were further selected for CD133. % of CD11b^+^’ CD11b^-^CD133^+^ and CD11b^-^CD133^-^ cells were calculated to initial cells seeded and plotted. (E) CD133^+^ cells and CD11b^-^ cells were further transfected once with miRs and stained for neuronal markers Nurr1 and Map2 and observed through immunofluorescence microscope. (400X magnification).

Presence of Prominin-1 indicated that miR may also be manipulating Yamanaka’s (OSKM) reprogramming factors. Presence of Prominin alone does not confirm the stemness of a cell as many epithelial mature cells express CD133 (Corbeil et al., 2014). To evaluate potential role of selected miR in manipulation of OSKM transcription factors, the cells after 2^nd^ transfection were analysed for the OSKM and other pluripotency genes. Core reprogramming factors *oct4, sox2, klf4, c-myc* and *nanog* mRNA expressions were all increased at various levels during macrophage reprogramming. *sox2* and *klf4* (Fig. 5B) expression increased in transfected cells correlating with increased prominin-1 (Fig. 5A) indicating that our strategy of macrophage reprogramming also involves manipulation of pluripotency. *sox2* is a transcription factor responsible for ectodermal transformation in iPSC (Johansson and Simonsson, 2010) and is expressed in cells transfected with all the miRs inducing neuronal differentiation. *c-myc*, an oncogene and *Pu.1*, myeliod lineage transcription factor, were decreased in all the transfected cells while an increased *oct4* was observed in miR 124 and miR 155 transfected cells (Fig. 5B). The reprogramming factors are reported to bind to miR 302 promotor and helps in its expression (Card et al., 2008). mRNA levels of miR 302, a known pluripotency inducing miR, was also increased following miR transfection indicating a possibility of miR-miR regulation as well (Fig. 5C). miR 302 has been shown to govern self-renewal and promote somatic cell reprogramming (Zhang et al., 2015). miR 224 transfection increased miR 367 levels (fold increase, 2.4±0.3) as analysed by real time PCR (data not shown) and miR 367/302 is shown to play a role in somatic cell reprogramming (Zhang et al., 2015). Although, an alteration in pluripotency genes was observed, but it does not indicate the key stem cell property as no cell division was detected after reprogramming protocol. Thus, these miR mediated cellular reprogramming, despite of manipulating the OKSM factor, does not results in iPSC but may have a transient stem-like phenotype. To further confirm the stemness, Prominin^+^ cells were purified after second transfection and were maintained in F-12 media (Fig. 5D). Despite being CD133^+^, these cells did not result in embryoid body formation or colonogenicity (the key features of iPSCs) after 7 days (data not shown).

The role of miR 9, miR 124 and miR 155 in differentiation and propagation of neural cells has been studied extensively. Many reports suggest that elevated levels of these miR ‘s in neurons. Thus, we explored the potential of these miRs in differentiating (Prominin) CD133^+^ cells obtained during the current reprogramming protocol. After the second transfection CD133^+^cells were isolated and re-transfected with miRs. Except miR 224, the other miRs were able to differentiate the CD133^+^ cells to neurons. This indicates that miR 9, 124 and 155 could reprogramme stem like-cells and trans-differentiate them to neurons while reprogramming with miR 224 follows direct reprogramming (Fig. 5E).

### miR-induced iNCs were functionally active

Synchronous network activity is integral for the development and functioning of brain. The balance of excitation of neurons relies on release of Ca^+2^ which further regulate expression of genes and release neurotransmitters at synaptic junction. Synapsin is functional protein present at the synaptic junctions indicating the functionality of neurons. It has been reported that Ca^+2^ plays vital role in driving differentiation and the frequency of Ca^+2^ transitions echoes various stages of neuronal maturation (Toth et al., 2016). To investigate the generic excitability of iNCs, intra cellular Ca^+2^ evoke was recorded at an interval of 30 secs. It was observed that the generated neurons demonstrate Ca^+2^ spikes following the change in the transmembrane ionic potential via addition extracellular potassium chloride. The Ca^+2^ immunoflouresnce intensity, through pseudo colour analysis (Fig. 6A, B_1-2_) correlated the recorded change in total fluorescence intensity (Fig 6C). Flow cytometry data also demonstrate that miR 9, miR 124 and miR 155 generated functionally active neurons (Fig. 6D and E and supplementary video SV1 and SV2). High expression of synapsin in generated iNCs by Western blotting (Fig. 6F and F’) and immunofluorescence staining (Fig. 6G and supplementary figure S5) indicate the possibility of inter-neuronal interactions and functionality.

**Fig. 6:**
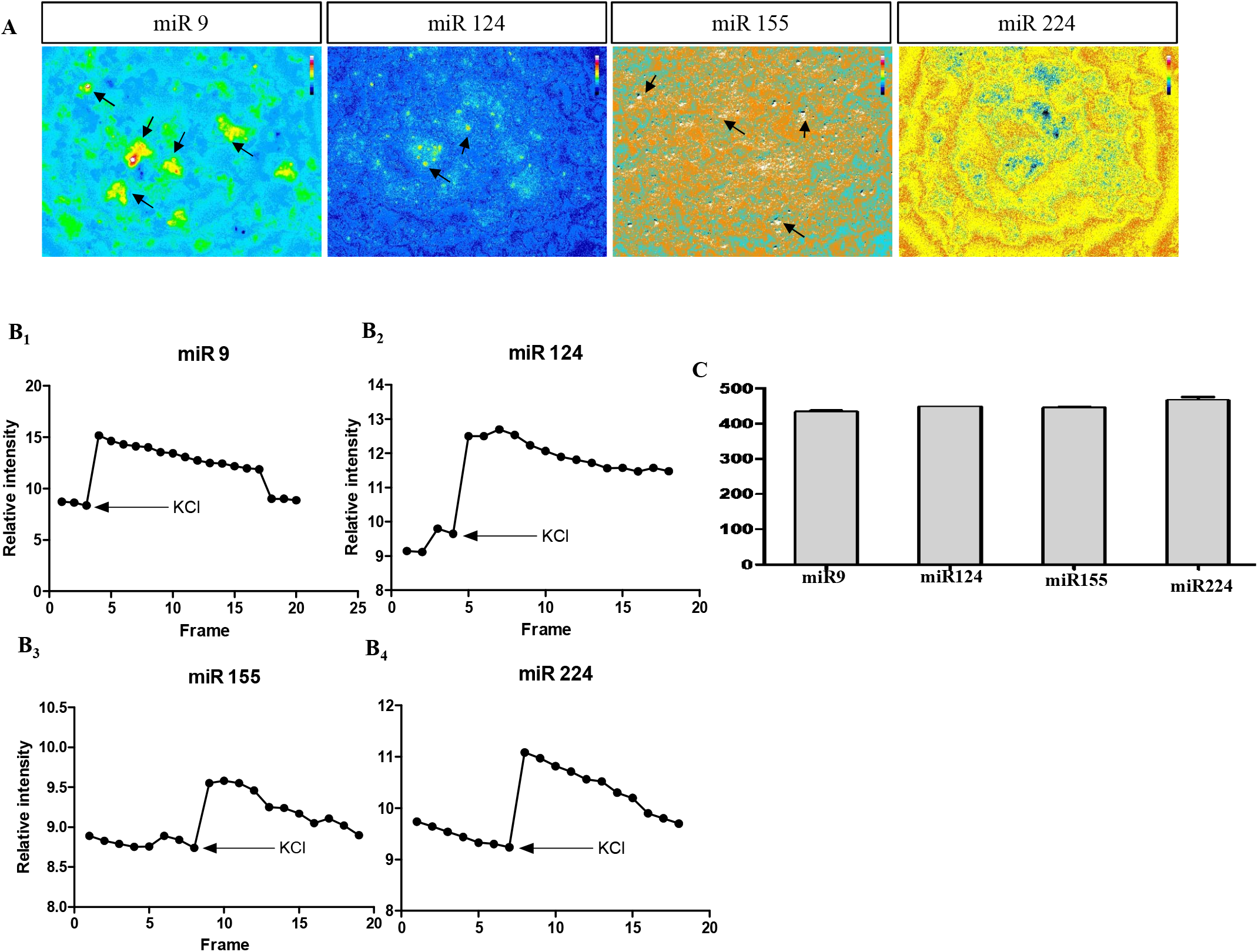
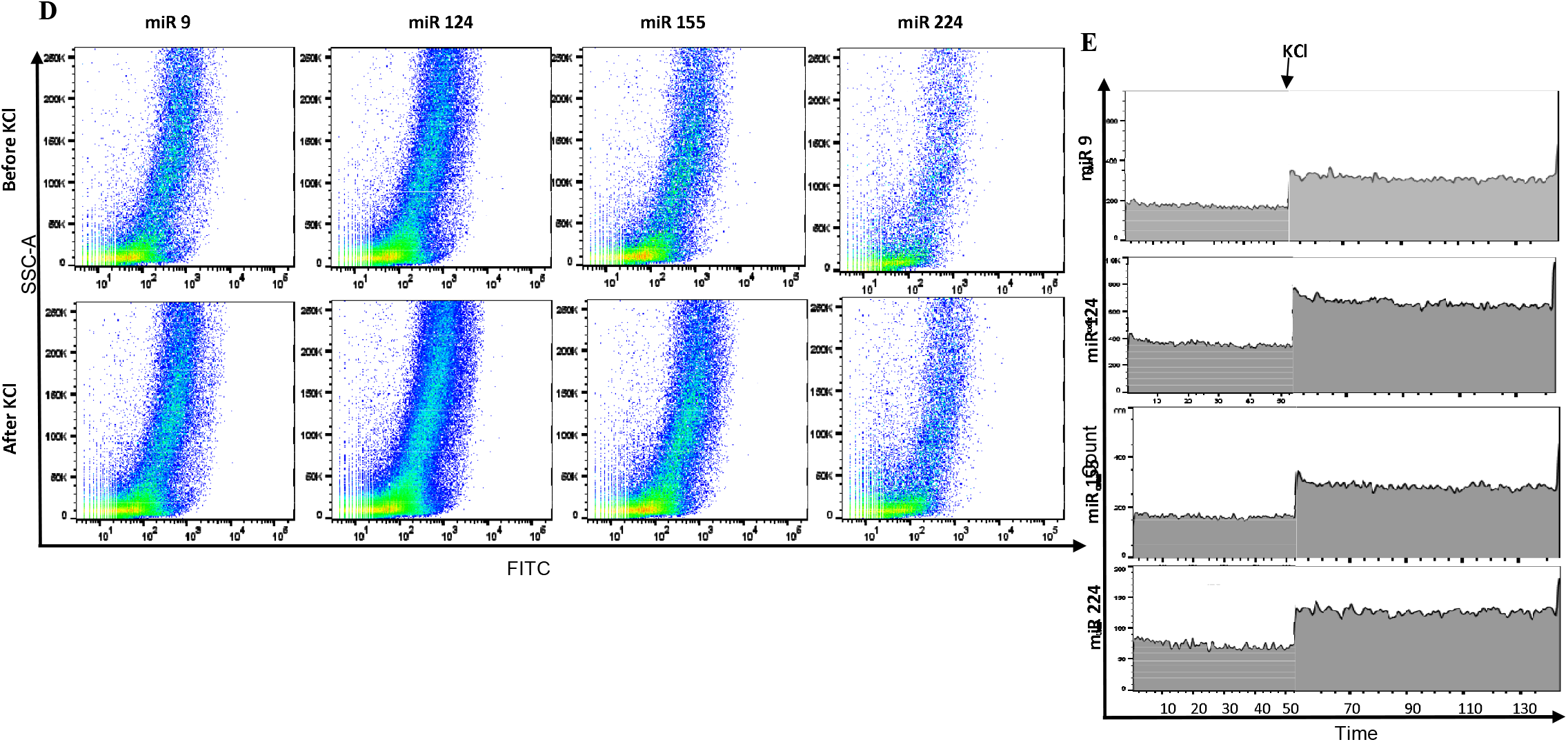
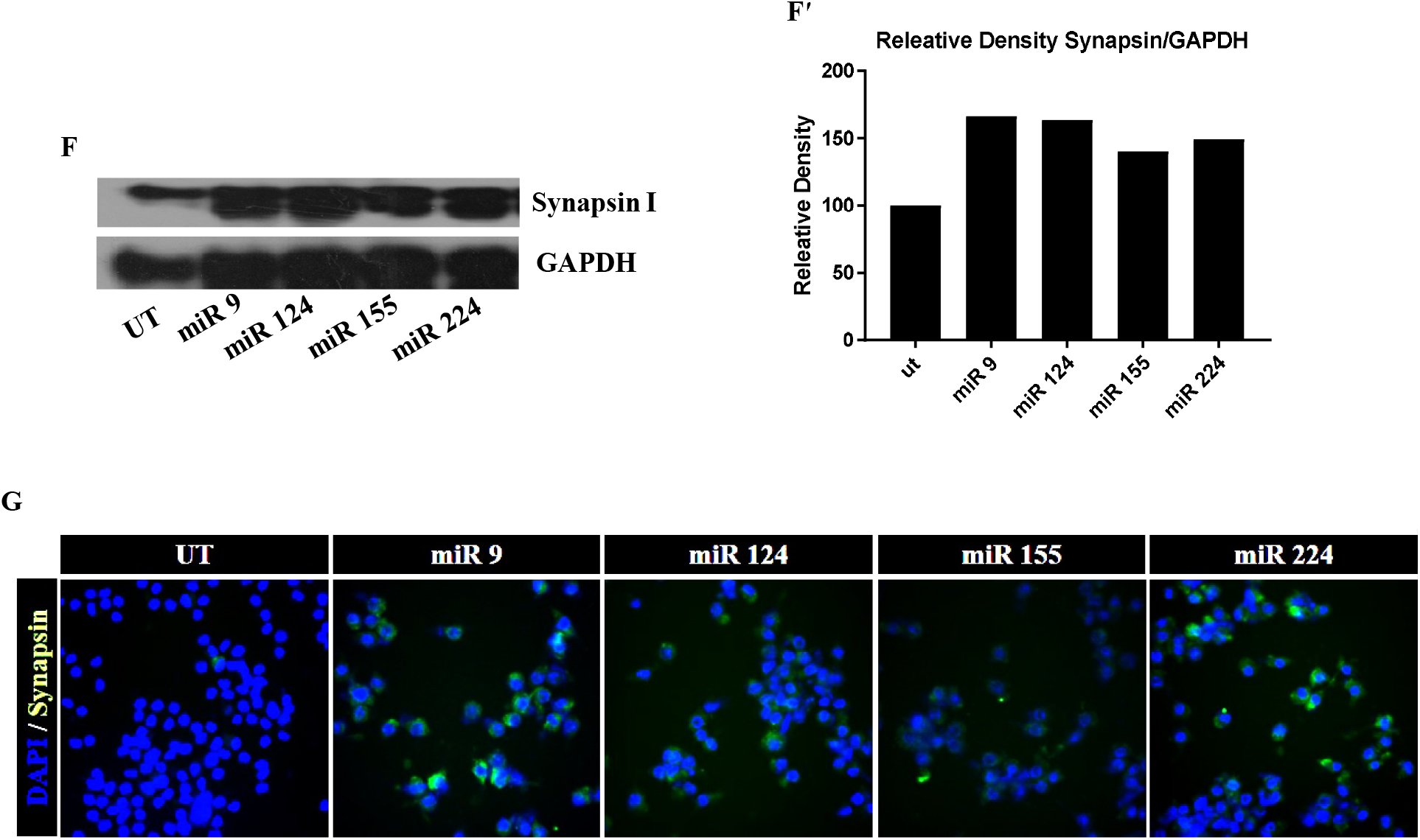
iNCs were physiologically active and express Synapsin I. Fluo-4, AM treated cells were imaged through period of time and merged together with pseudocolor to represent increase in intracellular calcium (A). Images were further analysed and increase in intensity was represented as Relative intensity vs Frame (B_1-2_). (C) Mean Fluorescence Intensity was measure before and after addition of KCl and plotted as change in MFI. iNCs were treated with Floua-4AM and analysed in flow cytometry green filter before and after addition of Kcl and plotted as (D) FITC vs SSC-A in pseudocolour. (E) FITC vs time. Synapsin is highly expressed in iNCs as analysed by Western blot (F and F’) and immune-fluorescence (G). (Fluo – 4, AM was captured at 200X magnification; immunofluorescence at 400X magnification). (see also Supplementary figure S5).

### Selected miRs can also induce reprogramming in mouse primary macrophages to iNCs

To test the efficacy of selected miR for reprograming primary microphages, we have used the bone marrow derived macrophages (BMDM) and peripheral blood mono-nuclear cells (PBMCs). BMDMs and PBMCs being negative for pluripotency genes exclude any possibility of stemness. The BMDMs were also negative for early and late neuronal markers. Transfection of BMDM’s with miR’s generated neurons with robust expression of Map2 and synapsin (Fig. 7A). Expression of NSE and N-Cadherin were also observed in generated iNCs (Fig. 7B). The protocol was confirmed by transdifferentiating PBMCs to neurons. miR transfection of PBMCs generated Map2+ neurons after three transfections (Fig. 7C), however miR 224 failed to generating neurons as it doesn’t have any role in neurogenesis (Choy et al., 2017).

**Fig. 7:**
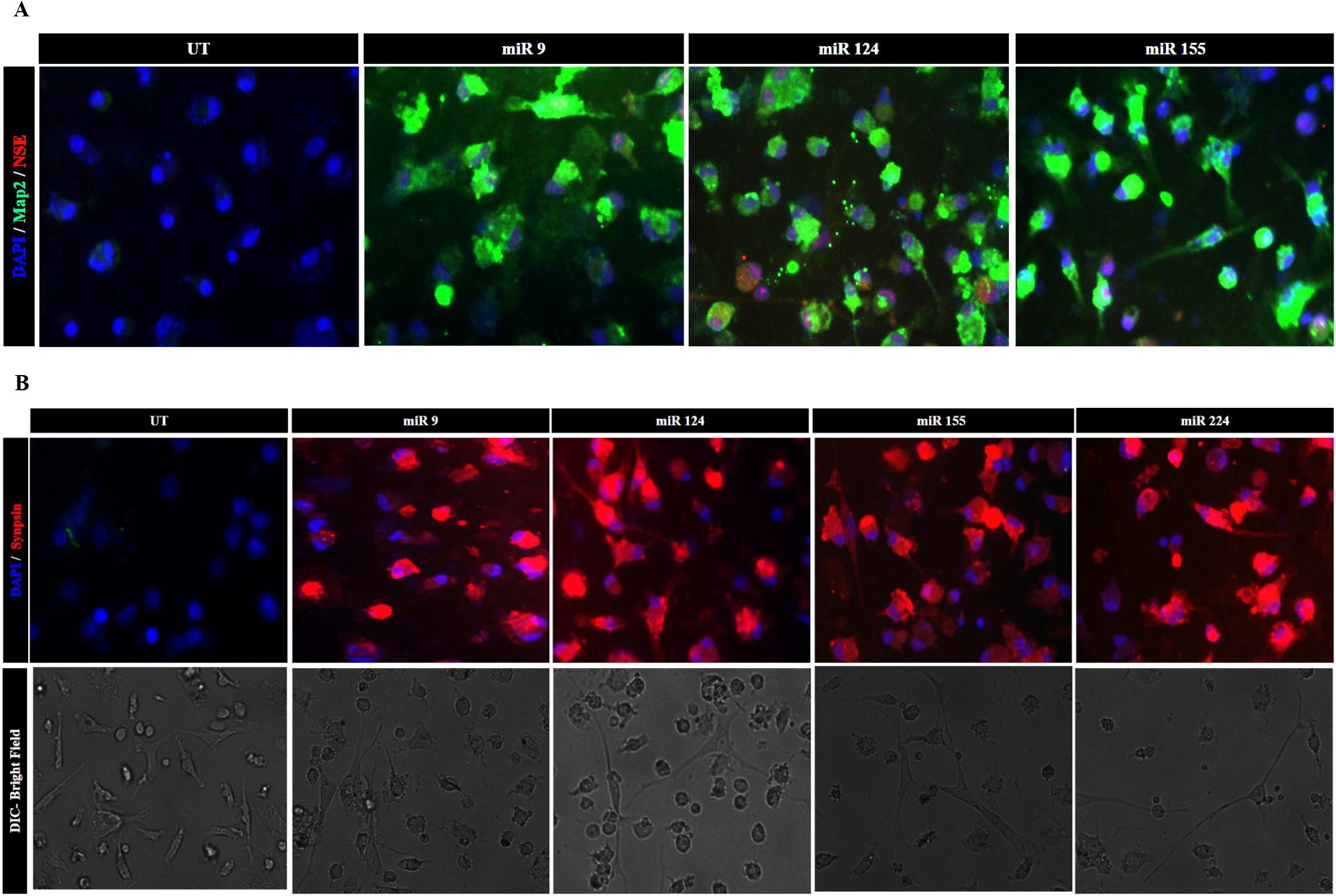
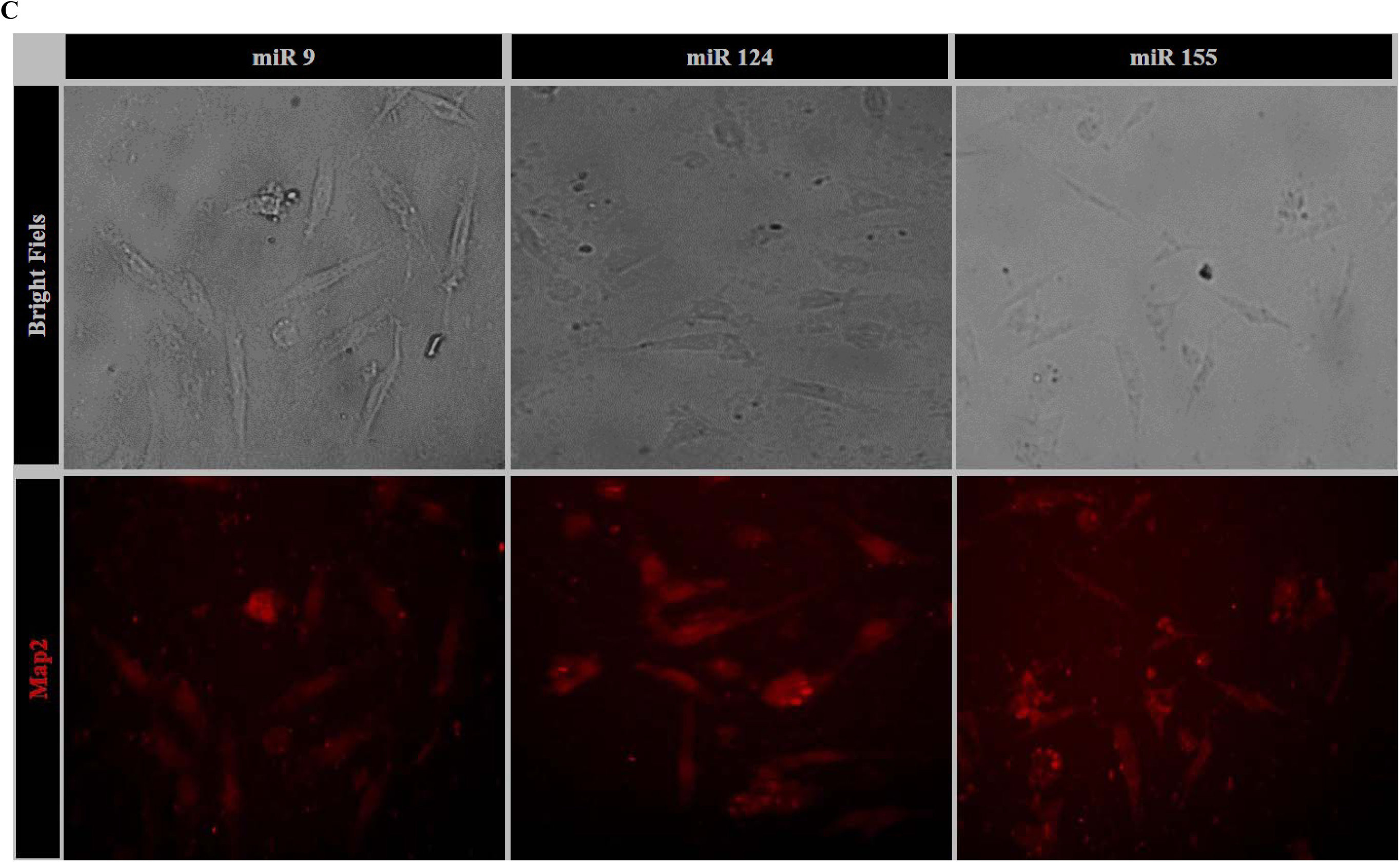
Mouse primary macrophages were transdifferentiated to iNCs. iNCs generated from bone marrow derived were analysed for Map2, NSE (A) and Synapsin (B). iNCs generated from PBMCs express Map2 as visualized by immunofluorescence (C). (400X magnification)

To validate the reprogramming potential of these miRs, we have also transfected fibroblasts, cells of different origin. The transfected fibroblast also attains neuronal like morphology with elongated neurites (supplementary figure S6) and expressed neuronal Markers Map2 and NSE (data not shown) after three transfections.

Overall we demonstrate high potential of selected miRs in cellular reprogramming. Each of them, however demonstrated variations in the associated mechanisms leaving opportunity for further exploration.

## DISCUSSION

Transcription factors occupy a key position in regulating gene function specificity in various cell types including many other functions. Yamanka and others have demonstrated transcription factor (OSKM) driven cellular reprogramming that established them as key fate determinants active during development. Their aim was to reprogramme somatic cells to various lineages different from their origin. miRNA-mediated repression regulates cell fate including its division and diversification. miRs are important post transcriptional repressors and regulate their target gene levels to confer diversity in a defined organ and its function making it an attractive target for cellular reprogramming. Their importance is also heighted by Hobertt et al., in 2008 which demonstrated that deletion of less than 10% miR caused broader defects compared to the deletion of 30% of the transcription factors. This was explained by the fact that the miRs expressions are much tightly regulated compared to the transcription factors. miR alterations govern many cellular processes and functions including haematopoiesis (miR17, miR 150, miR 155, miR221, miR222, miR 424) (Bissels et al., 2012), cell division (miR193) (Pruikkonen and Kallio, 2017) and pluripotency (miR 302/367) (Zhang et al., 2015). The current study aimed at trans differentiation of macrophages to neurons. Owing to the non-replicative nature of neuron it is desirable to transdifferentiate other cell types to functional neurons for experimental and clinical applications. Over the years, direct reprogramming has proven that cells from various lineages can be converted to cells of alternative linages, even across the germ layers. Lineage boundaries can thus be overcome, depending on the potency of the factors employed (Masserdotti et al., 2016). Stem cells of mesenchymal origin, skin fibroblasts, astrocytes, hematopoietic stem cells, urinary cells and adipocyte progenitor cells were successfully transformed to neurons using various reprogramming strategies (Biswas and Jiang, 2016). However, cells of ectodermal origin form an attractive target owing to the ease of conversion to neurons, it involves but it is an invasive procedure. The mesenchymal stem cells are easy to reprogramme, but they have their own ethical concerns. Despite of being terminally differentiated macrophages they demonstrate structural and functional plasticity in various tissues (Galli et al., 2011). However, their conversions to other cell type remains unexplored. Hence, in the current study we choose to reprogramme macrophages owing to interesting factors like ease of availability in the host, cellular excitability, presence of shared signalling molecules and extensive cross talk with the nervous system. Previously, JH Lee at al., reprogrammed blood progenitors from cord blood to neuronal progenitors using *Oct-4* overexpression and small molecules, as the blood cells can be easily obtained, purified and stored (Lee et al., 2015). Overexpression of neurogenin 2 and co-culture with neural cells confer neurogenic potential to monocytes, which also lead us to opt for macrophages (Kodama et al., 2006). Multiple targets manipulated by miR provide a wider and acceptable changes in the cell fate without the possibility of oncogenic transformations. We have selected screened miRs which are expressed during different development stages of neurons and their essentially have been established for diverse functions including cellular morphology in neurons. Nonetheless, miR expression vectors carrying miR9-124-Bcl was sufficient to reprogramme fibroblasts to neurons where Bcl manipulated apoptosis while miR9 and miR124 served to direct their conversion to neurons (Pfisterer et al., 2016). The exclusive use of miRs for direct reprogramming has not been reported till date. Macrophages and neurons have different development origin. The miRs were judiciously selected taking into account the ability to differentiate the cells beyond the boundaries of tri-germinal layer and are expressed during different developmental stages of neurons. We demonstrated that enrichment of miRs via mimics can exclusively generate iNCs without the use of Yamanaka factors, cell cycle manipulators or small synthetic molecules. *In silico* analysis, however predicts that the miRs used in the study may manipulate pluripotency factors via direct or indirect interactions.

In the current study, we aimed to utilize miR-assisted differentiation of macrophages for generating functional neurons. We showed that macrophages transfected with miR 9, miR 124, miR 155 and miR 224 transdifferentiated to neuron-like cells in neuronal induction media (NIM). The NIM included: Iso-butylmethyl Xanthine (IBMX), Neural growth factor (NGF) and Retinoic Acid (RA). IBMX is known to manipulate protein kinase A (PKA) pathway, a crucial mediator for neuronal differentiation and enhances neurite length (Tio et al., 2010). RA plays a crucial role in differentiation by binding to RA receptors and modulating Wnt signalling a known pathway in neuronal differentiation. NGF is a growth factor promoting neuronal differentiation by inducing Tubb3, and Nurr1 expression and is involved in bringing the biochemical and electrophysiological changes that recapitulate certain features of sympathetic neurons (Lee et al., 2013). Tio et al., differentiated mesenchymal stem cells to neurons using IBMX, RA, FGF and NGF (Tio et al., 2010). NIM along with miRs derived neuronal gene expression in macrophages may be transdifferentiating them to neurons.

miR 9 has been demonstrated to induce neurogenesis, but has never been explored for cellular reprogramming. Reports suggest that miR 9 differentiates neural stem cells (NSCs) by downregulating Nuclear Receptor Subfamily 2 Group E Member 1 (NR2E1, TLX) (Radhakrishnan and Alwin Prem Anand, 2016). It can also manipulate Wnt signalling that plays a major role in generation of dopaminergic neurons and REST expression that forms a feedback loop to control neuronal differentiation. String 10.0 analysis of miR 9 predicted target genes show involvement of functional neurogenic genes (Fig S1). Furthermore, pak4, scrib and prrx1 that regulate klf 4 expression via Rho proteins were predicted to be manipulated by miR 9 as evidenced by increase in klf 4 expression by miR 9 mimic (Fig. 4B, 5A). The *in silico* analysis showed interaction with Foxg1, a validated target for miR 9, which plays a role in differentiating neurons during early stages of embryonic development (Yuva-Aydemir et al., 2011). miR 9 target, sirt1 may be the driving factor for neuronal differentiation (Liu et al., 2014) (Fig. 1C) which is concurrent with a recent report (Saunders et al., 2010). *In silico* analysis also predicted inhibition of REST1, a well-known target of the miR −9/9*, that has an important role in neurogenesis. It is thus perceived that miR 9 might inhibit Wnt/catenin signalling pathways (Yu et al., 2014) thus enhancing reprogramming ability as demonstrated (Ho et al., 2013). In aggregate, miR 9 mediated neurogenesis can be explained via inhibition of like GSK3-β, REST1, SIRT1, Wnt/catenin and other possible targets which may be aiding to initiate and promote reprogramming of macrophages into neurons.

miR 124 has earlier been shown to promote neuronal gene expression suggesting its pro-neural role (Victor et al., 2014). The pro-neural functions of miR 124 have been attributed to at least these five mechanisms: i) REST/small CTD phosphatase 1 pathway ii) Inhibition of BAF complex iii) Suppression of sox9 expression iv) Down regulation of PTBP1 v) regulation of ephrin-B1 mRNA stability. Among all the tested miRs, we demonstrate that miR 124 possessed a remarkable ability to transdifferentiate macrophages into neurons with an efficacy of 18% (Fig. 3). The mechanisms mentioned earlier may also to be involved in reprogramming based on predictions by bio-informatics analysis. Report by, Xio et al., in 2015 indicated that the decreased FOXA2 expression following miR 124 inhibition is mandatory for maintenance and development of notochord and neural tube (Dal-Pra et al., 2011; Masserdotti et al., 2016). miR 124 mimic may be either restoring or increasing FoxA2 levels which may facilitate the expression of neuronal genes. Interaction of miR 124 with Sox9 may be involved in mediating neuronal differentiation. Surprisingly, *in silico* analysis also picked up Jagged1 ligand of notch signalling (Fig. 4C), suggesting that its inhibition by miR 124 may play a role in reprogramming (Ichida et al., 2014). Neuronal differentiation by miR 124 mimic (Fig. 5E) can thus be partly explained by intermittent expression of pluripotency and neuronal genes. We thus perceived that trans differentiation of macrophages into neurons could be explained by the above mechanisms of miR 124 targeted genes as depicted in string analysis (Fig. 5C) regulating pluripotency and neuronal genes.

miR 155 has an important role in early developmental and its expression is increased in blastocysts indicating its function in self-renewal (Goossens et al., 2013). TGF-β, an important regulator of cellular plasticity and cell fate also manipulates miR 155 expression thus indicating its involvement in cellular reprogramming (Kong et al., 2008). miR 155 plausibly inhibits the myeloid lineage maintenance in macrophages, via decrease in *PU.1* (Table S3, Fig. 2B, 2B’ and 5B) and components of the NIM may instigate reprogramming to functional neurons. I*n silico* analysis predicted Hif1, Foxa2, Nrf2 and sox2 as the targets of miR 155 which are also reported to be involved in reprogramming (Fig. S1). The manuscript demonstrates that in addition to the potentiating neurogenic genes (Fig. 2 and 3), miR 155 mediated reprogramming to iNCs is via inhibition of the macrophage lineage. miR 155 transfection upregulated pluripotency genes like *oct4* and *nanog* and drive cellular reprogramming (Fig. 5B). Even though there is no report claiming neurogenic potential of miR 155, we for the first time demonstrate the neurogenic potential of miR 155 with reprogramming efficacy of 21% (Fig. 3B). The mechanism of cellular reprogramming may thus be facilitated via dedifferentiation of macrophages and NIM assisted neurogenesis rather than direct trans-differentiation.

miR224 also trans-differentiated macrophages into neurons, but the efficacy was least compared to other miRs used in the study. miR 224 manipulates Wnt/catenin signalling pathway and PDGFRA, which are involved in regulating stemness and pluripotency during early developmental events (Yu et al., 2014; Zhu et al., 2014). Foxm1, RuvB-like protein 2, Atf6b, Spag1, Ndrg3 showed up during *in silico* analysis (Fig S1) which are also reported for maintenance and proliferation of embryonal stem cells and neuro-ectodermal differentiation (Do et al., 2014; Kwok et al., 2016). Concurrently, our results demonstrate that miR 224 could transform macrophages into neurons via cells expressing pluripotency genes (Fig. 5B). However, the low turnover (Fig. 3B) could be attributed to insufficiency of miR 224 in inducing neurogenic genes or supressing myeloid lineage maintenance genes.

The novel strategy of miR-induced trans differentiation of macrophages to neurons offers a technological advancement over other reported protocols with respect to the ease of cell availability, robustness, non-invasiveness and less time consuming and better turnover with respect to the quantity (percent of cell recovery) and quality (synapsin^+^ excitable) neurons. The study further warrant additional information regarding the molecular mechanism of cellular reprogramming and also requested to its pre-clinical and clinical efficacy. miR-mediated reprogramming is more feasible for clinical applications, *in vivo* reprogramming, site directed reprogramming.

## MATERIALS AND METHODS

### Cell lines

The mouse macrophage cell line, Raw 264.7 and mouse fibroblast cell line, L929 were procured from National Centre for Cell Science (NCCS, Pune, India) and used. The cells were cultured and maintained in Dulbecco’s Modified Eagle’s Media (DMEM) (Thermo Fischer Scientific, USA) supplemented with 10% Fetal Bovine Serum (FBS) (GIBCO), 1X Pencillin-streptomycin-neomycin (Thermo Fisher Scientific, CA, USA). The cultures were maintained at 37°C in a humidified incubator with 5% CO_2_. All the miR mimics were purchased from Qiagen, Hilden, Germany. Isobutylmethylxanthine (IBMX) and Retinoic Acid (RA) were procured from Sigma Aldrich, Missouri, USA; and Nerve Growth Factor 7S was purchased from Thermo Fisher Scientific, CA, USA. All the reagents were procured from Thermo Fisher Scientific, USA until unless mentioned.

### Primary cultures

Bone marrow cells were collected from BALB/c mice. The marrow was collected from Femur and single cell suspension was prepared in RMPI with 10% FBS, sodium pyruvate and HEPES. Cells were maintained in monocyte colony stimulating factor (14-8983; eBiosciences; 30 ng/ml) as described earlier (Gaikwad et al., 2017) for 7 days for macrophage formation. Adherent cells were washed and transfected for reprogramming. Peripheral blood was collected by retro-orbital bleeding from BALB/c mice. Peripheral blood mono-nuclear cells (PBMCs) were purified using Ficoll-gradient centrifugation. Cells were plated and adherent cells were washed before transfection for reprogramming.

Mouse miR 9, miR 124, miR 155 and miR 224 (Qiagen, HL, Germany) (with sequences as given in supplementary Table S1) were prepared in molecular biology grade water and the final concentration used for transfection was 5nM. miRs were transfected using Lipofectamine *RNAimax* in serum free Opti-MEM for 24 h, followed by maintaining cells in Neuronal Induction Media (NIM) for 48hrs or till further transfection (Fig. 1A). NIM was prepared to final concentrations of 510μM IBMX, 50nM NGF and 10μM Retinoic Acid (in DMSO) in DMEM + F-12 media containing 5%FBS. To check the transfection efficiency, the cells were collected after 48hr. For carrying out the functional analysis the cells were collected after the completion of the protocol as described in Fig. 1A.

### Real time-PCR

Transfection efficiency and altered gene expression were analysed using SBYR Green Real-Time PCR Master Mix (A25741; Applied biosystems). Briefly total RNA was extracted from cells using TRIzol (Thermo Fisher Scientific, CA, USA) as per manufacturer’s protocol. 1μg of total RNA of each sample was used for cDNA synthesis using RT-PCR System (Bio-Rad, USA) under following conditions: 80°C for 5 min, 60°C for 5 min, 60°C for 45 mins and 85°C for 5 min. The cDNA template was used for amplifying specific genes and miRs using gene and miR specific primers (Supplementary table 2) in Power SYBR green 2X Master Mix in Thermal cycler and analysed with SDS2.4 software (Applied Biosystems, USA). Following were the conditions used: 40 cycles followed by a final extension at 72°C for 10 min. Relative gene expression was normalized to internal control, GAPDH.

### Semi quantitative RT-PCR

Gene expression was analysed using semi quantitative RT-PCR as described earlier (Patel et al., 2018) using gene specific primers (Supplementary Table.3) Briefly, total RNA was extracted from cells using RNAiso (Takara Bio, USA) according to manufactures protocol. cDNA was synthesised as mentioned in previous section. Synthesised cDNA was used for analysing the expression of specific genes using gene specific primers (as given in supplementary table 1) in presence Taq polymerase master mix (EmaraldAmp GT PCR Master mix, Takara, Japan).

### Immunostaining

Neuronal markers were detected after treatments under immunofluorescence microscope (Olympus, Japan) at 40X objective as describe previously (Naveen et al., 2016). Briefly, transdifferentiated cells were fixed with 2% paraformaldehyde and washed with PBS. Cells were permibialised using 1.5% tween in PBS. Blocking was done with 2%FBS in PBS after washing thrice. Anti-Map2 a,b,c (MA5-12826; Thermo Fisher Scinetific;1:500), anti-Neuron Specific Enolase (NSE; PA5-12374; Thermo Fisher Scientific; 1:500), anti-nestin (MA1-110; Thermo Fisher Scientific; 1:500), anti-nurr1 (PA5-22799; Thermo Fisher Scientific; 1:500), anti-synapsin I (S193; Sigma;1:1000), anti-Prominin (SAB4300882; Sigma-Aldrich; 1:1000) (Sigma Aldrich, USA), anti-N-Cadherin (C3865; Sigma;1:500), anti-glial fibrillary acidic protein (GFAP; 14-9892; eBioscience, 1:1000) FITC-Labelled anti-CD11c (553801; BD biosciences; 1:500) were used as primary antibodies. Secondary antibodies like goat anti-mouse IgG/IgM Alexa flour 488/543 (A11029; Thermo Fisher Scinetific; 1:2000), goat anti-Rabbit Alexa Flour 546 (A11003; Thermo Fisher Scinetific; 1:2000) were used. All the images

### Western blotting

Expression of proteins were assayed by Western blotting as described earlier (Gaikwad et al., 2017). Briefly, cell lysates (100μg) were prepared in RIPA buffer. Proteins were separated in 10%SDS-polyacrylamide gels, and the protein were electro blotted on to nitrocellulose membrane. The membrane was probed with indicated primary of dilution 1: 1000 for overnight at 4°C and then incubated with respective HRP-labelled secondary antibodies [(anti-Rabbit IgG; 656120; Thermo Fisher Scientific), (anti-Mouse IgG; HP06; GeNei)] of dilution 1:3000 for 1 hr. GAPDH was detected as internal control using Anti-GAPDH (AM4300; Thermo Fisher Scientific, 1:1000). Secondary antibodies were detected by exposure to X-ray film after incubation with Clarity western ECL substrate (Bio-Rad). Images were represented of three individual experiments.

### Cell separation

After treatments, the cells were washed with washing buffer (PBS containing 1%FBS) and incubated with biotin labelled anti-CD11b antibody (RM2815; Invitrogen; 1:100) for 40min with constant mixing using Hula mixer (Invitrogen) at room temperature. Excess antibody was washed form the cells by centrifugation at 600 g for 2 min. After 2 washes the cells were incubated with Dynabeads MyOne streptavidin C1 for 20 mins at room temperature. CD11b^-^ and CD11b^+^ cells after magnetic separation were re-suspended in DMEM and washing buffer respectively for further analysis. CD11b^-^ cells were further enriched for CD133^+^ and CD133^-^ cells. Purified cells were counted and plated on poly-Ornithine (Sigma Aldrich, USA) coated plates. Experimentation was carried out as mentioned in respective sections.

### Flow cytometry

Cells were collected after each transfection and maintained in NIM. Flow cytometry was done for neuronal marker expression by as described earlier (Patel et al., 2018). Briefly, cells were collected in ice cold PBS and washed in FACS buffer and fixed in 2% paraformaldehyde. For MAP2 and NSE staining, cells were permibialised using 2% Tween-20 in PBS. Primary antibodies anti-mouse MAP2 (1:400), anti-mouse NSE (1:500), anti-Synapsin (1:500) and anti-CD133 (1:500) were used after Fc blocking with CD16/32 antibody (553141; BD Pharmingen). Respective Secondary antibodies: goat anti-mouse IgG/IgM Alexa flour 488 (1:1000), goat anti-Rabbit Alexa flour 546 (1:1000) were used. Samples were acquired in BD FACS Canto II and analysed using BD Cell Quest Pro software. Efficacy was calculated as % of neuronal cells obtained to macrophages seeded (formula given in supplementary data).

### Intracellular calcium measurement

Intracellular calcium was measured in three different ways in this paper using Fluo-4, AM dye (F14201; Thermo fisher, USA) which specifically binds to intracellular calcium. Cells were treated with Fluro-4AM dye for 30 mins in HBSS buffer, washed and analysed within 30 mins. For flow cytometry, cells were acquired in FACS canto where no data acquisition was recorder for first 10 sec. After 10 Sec of acquisition without acquiring data, and acquired the data for next 10sec, 1mM potassium chloride (KCl) was added and continued acquiring the data immediately after adding activation for 30sec. Result was shown in SSC vs MFI (Fig. 6D) and as intensity vs time (Fig. 6E). For immuoflourescence analysis the cells were observed in Olympus, Japan at Excitation/ Emission: 488/520nm. After addition of activator, images were captured every 2sec for 1 mins continuously. All the images of were stacked and merged together and processed with macro tool - dFoverFmovie of Image J, NIH, USA. 16_colour LUT was applied to colourise the images. To further quantify the Fluo-4, AM intensity, the live differentiated iNCs were cultured in 96 well plates in triplicates.

Fluorescence was measured in multimode reader (Molecular devices, USA) in before and after the addition of KCl. The represented graphs are normalized as increase in mean fluorescence intensity.

### *In silico* miR target prediction and functional co-relation

To select brain/neuron specific miRs, the expression profiles of various miRs were screened. Tissue specific miRs screening was done using miRGator v3.0, EMBL-EMI, mESAdb and miRTargetLink that majorly based on transcriptome datasets. miR predicted targets were obtained using TargetScan 7.0 and miRGator and were selected for further analysis. Top 100 genes from TargetScan 7.0 and overlapping genes from miRGator (as given in Fig. S1) were considered for the further analysis. Experimentally validated miR targets were obtained from miRecords and were also added as input to the above list. Pool of miR target genes from all the three tools as given in the supplementary table no. 3 were given as input for gene ontology and function prediction. Pathways were enriched in PANTHER 11 and FAME using gene ontology (GO). STRING 10.0 analysis of the genes obtained in TargetScan was used to depict the molecular interaction between miR predicted targets and reprogramming factors and later functional ontology was applied and visualised in cytoscape.

The animal experiments were approved by the Institutional Animal Ethics Committee (IAEC) and Committee for the Purpose of Control and Supervision of Experiments on Animals (CPCSEA) at the B.V. Patel PERD Centre, Ahmedabad, Gujarat, India.

### Statistical Analysis

All the data are representative of minimum of triplicate data generated from a minimum of three individual experiments. All data are expressed as mean±S.D. P values were calculated using students’ two tailed paired t-test or one-way ANOVA, and a P<0.05 was considered statistically significant.

## Abbreviations

BMDM: Bone marrow derived macrophages
iNCs: Induced neuronal cells
iPSC: Induced pluripotent stem cells
miR: microRNA
NIM: Neuronal Induction Media
OSKM: Oct, Sox, Klf and Myc

## ACKNOWLEDGMENT

Authors acknowledge Mr. Manthan Patel and Ms. Kshama Jain for initial standardization of transfection and NIM the protocols; and Dr. Vinoth Kumar K for inducting qPCR technique. We thank Manish Nivsarkar for extending help in animal experiments. We thank ISSCR-CiRA 2017 and Immunocon 2017 (India) conference delegates for valuable suggestions. We thank Dr. Dhaval Patel and Dr. Suvendu Das for helping for suggestion in drafting the manuscript.

## AUTHORS CONTRIBUTION

RAR conceived the study. RAR and NC designed the experiments. NC performed the experiments. NC and RAR prepared manuscript.

