## Supplemental Figure for "microRNAs (miR 9, 124, 155 and 224) transdifferentiate macrophages to neurons"

### Supplementary File

#### microRNAs (miR 9, 124, 155 and 224) direct reprogramming of macrophages to neurons without the use of exogenous transcription factors

Naveen Challagundla[1], and Reena Agrawal-Rajput[1*]

[1]Immunology Lab, Indian Institute of Advanced Research [IIAR], Gandhinagar, Gujarat. 382427 India

**Figure S1**

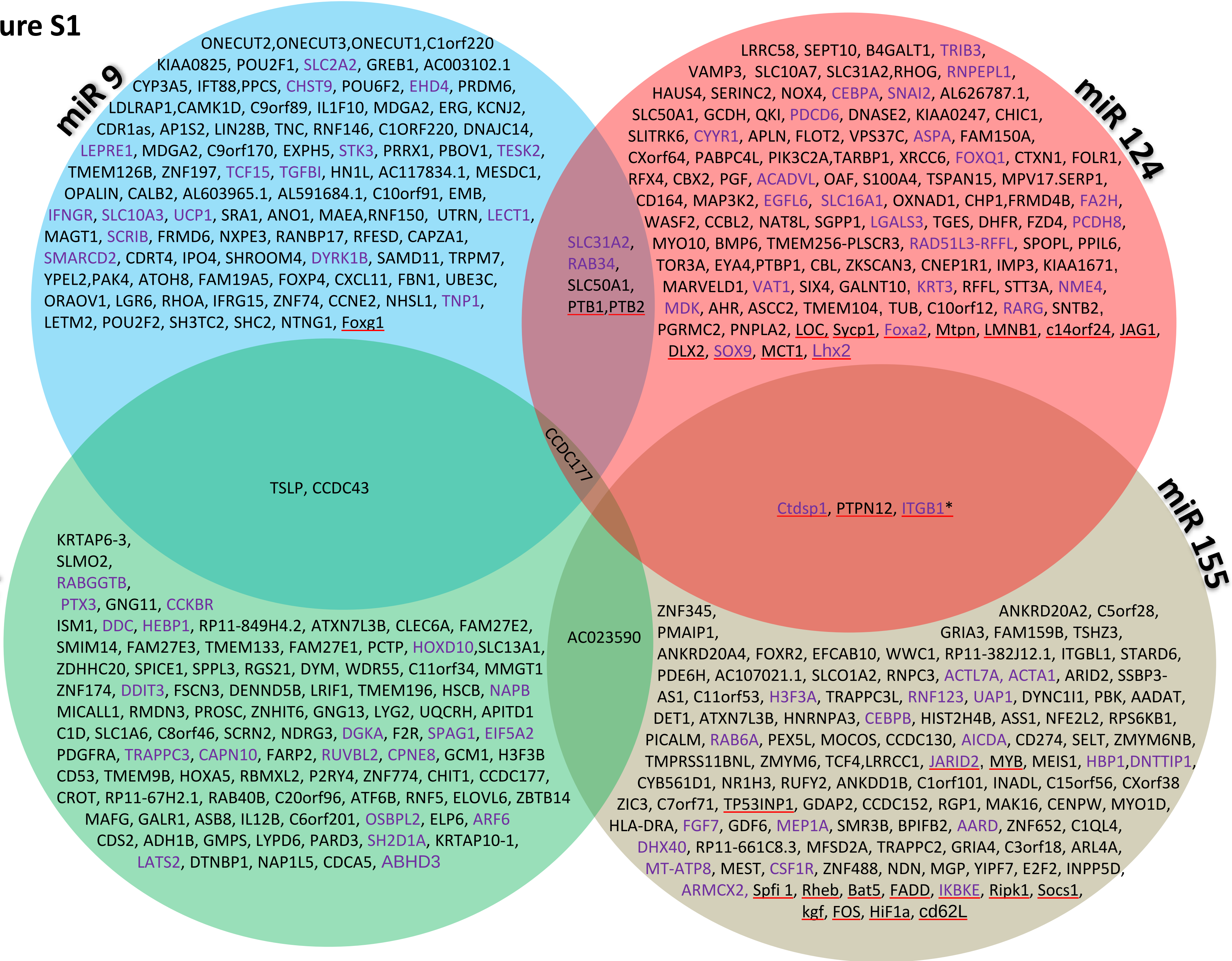

miR 9
ONECUT2,ONECUT3,ONECUT1,C1orf220
KIAA0825, POU2F1, SLC2A2, GREB1, AC003102.1
CYP3A5, IFT88,PPCS, CHST9, POU6F2, EHD4, PRDM6,
LDLRAP1,CAMK1D, C9orf89, IL1F10, MDGA2, ERG, KCNJ2,
CDR1as, AP1S2, LIN28B, TNC, RNF146, C1ORF220, DNAJC14,
LEPRE1, MDGA2, C9orf170, EXPH5, STK3, PRRX1, PBOV1, TESK2,
TMEM126B, ZNF197, TCF15, TGFBI, HN1L, AC117834.1, MESDC1,
OPALIN, CALB2, AL603965.1, AL591684.1, C10orf91, EMB,
IFNGR, SLC10A3, UCP1, SRA1, ANO1, MAEA,RNF150, UTRN, LECT1,
MAGT1, SCRIB, FRMD6, NXPE3, RANBP17, RFESD, CAPZA1,
SMARCD2, CDRT4, IPO4, SHROOM4, DYRK1B, SAMD11, TRPM7,
YPEL2,PAK4, ATOH8, FAM19A5, FOXP4, CXCL11, FBN1, UBE3C,
ORAOV1, LGR6, RHOA, IFRG15, ZNF74, CCNE2, NHSL1, TNP1,
LETM2, POU2F2, SH3TC2, SHC2, NTNG1, Foxg1
miR 124
LRRC58, SEPT10, B4GALT1, TRIB3,
VAMP3, SLC10A7, SLC31A2,RHOG, RNPEPL1,
HAUS4, SERINC2, NOX4, CEBPA, SNAI2, AL626787.1,
SLC50A1, GCDH, QKI, PDCD6, DNASE2, KIAA0247, CHIC1,
SLITRK6, CYYR1, APLN, FLOT2, VPS37C, ASPA, FAM150A,
CXorf64, PABPC4L, PIK3C2A,TARBP1, XRCC6, FOXQ1, CTXN1, FOLR1,
RFX4, CBX2, PGF, ACADVL, OAF, S100A4, TSPAN15, MPV17.SERP1,
CD164, MAP3K2, EGFL6, SLC16A1, OXNAD1, CHP1,FRMD4B, FA2H,
WASF2, CCBL2, NAT8L, SGPP1, LGALS3, TGES, DHFR, FZD4, PCDH8,
MYO10, BMP6, TMEM256-PLSCR3, RAD51L3-RFFL, SPOPL, PPIL6,
TOR3A, EYA4,PTBP1, CBL, ZKSCAN3, CNEP1R1, IMP3, KIAA1671,
MARVELD1, VAT1, SIX4, GALNT10, KRT3, RFFL, STT3A, NME4,
MDK, AHR, ASCC2, TMEM104, TUB, C10orf12, RARG, SNTB2,
PGRMC2, PNPLA2, LOC, Sycp1, Foxa2, Mtpn, LMNB1, c14orf24, JAG1,
DLX2, SOX9, MCT1, Lhx2
SLC31A2,
RAB34,
SLC50A1,
PTB1,PTB2
CCDC177
TSLP, CCDC43
Ctdsp1, PTPN12, ITGB1*
miR 224
KRTAP6-3,
SLMO2,
RABGGTB,
PTX3, GNG11, CCKBR
ISM1, DDC, HEBP1, RP11-849H4.2, ATXN7L3B, CLEC6A, FAM27E2,
SMIM14, FAM27E3, TMEM133, FAM27E1, PCTP, HOXD10,SLC13A1,
ZDHHC20, SPICE1, SPPL3, RGS21, DYM, WDR55, C11orf34, MMGT1
ZNF174, DDIT3, FSCN3, DENND5B, LRIF1, TMEM196, HSCB, NAPB
MICALL1, RMDN3, PROSC, ZNHIT6, GNG13, LYG2, UQCRH, APITD1
C1D, SLC1A6, C8orf46, SCRN2, NDRG3, DGKA, F2R, SPAG1, EIF5A2
PDGFRA, TRAPPC3, CAPN10, FARP2, RUVBL2, CPNE8, GCM1, H3F3B
CD53, TMEM9B, HOXA5, RBMXL2, P2RY4, ZNF774, CHIT1, CCDC177,
CROT, RP11-67H2.1, RAB40B, C20orf96, ATF6B, RNF5, ELOVL6, ZBTB14
MAFG, GALR1, ASB8, IL12B, C6orf201, OSBPL2, ELP6, ARF6
CDS2, ADH1B, GMPS, LYPD6, PARD3, SH2D1A, KRTAP10-1,
LATS2, DTNBP1, NAP1L5, CDCA5, ABHD3
AC023590
miR 155
ZNF345, ANKRD20A2, C5orf28,
PMAIP1, GRIA3, FAM159B, TSHZ3,
ANKRD20A4, FOXR2, EFCAB10, WWC1, RP11-382J12.1, ITGBL1, STARD6,
PDE6H, AC107021.1, SLCO1A2, RNPC3, ACTL7A, ACTA1, ARID2, SSBP3-
AS1, C11orf53, H3F3A, TRAPPC3L, RNF123, UAP1, DYNC1I1, PBK, AADAT,
DET1, ATXN7L3B, HNRNPA3, CEBPB, HIST2H4B, ASS1, NFE2L2, RPS6KB1,
PICALM, RAB6A, PEX5L, MOCOS, CCDC130, AICDA, CD274, SELT, ZMYM6NB,
TMPRSS11BNL, ZMYM6, TCF4,LRRCC1, JARID2, MYB, MEIS1, HBP1,DNTTIP1,
CYB561D1, NR1H3, RUFY2, ANKDD1B, C1orf101, INADL, C15orf56, CXorf38
ZIC3, C7orf71, TP53INP1, GDAP2, CCDC152, RGP1, MAK16, CENPW, MYO1D,
HLA-DRA, FGF7, GDF6, MEP1A, SMR3B, BPIFB2, AARD, ZNF652, C1QL4,
DHX40, RP11-661C8.3, MFSD2A, TRAPPC2, GRIA4, C3orf18, ARL4A,
MT-ATP8, MEST, CSF1R, ZNF488, NDN, MGP, YIPF7, E2F2, INPP5D,
ARMCX2, Spfi 1, Rheb, Bat5, FADD, IKBKE, Ripk1, Socs1,
kgf, FOS, HiF1a, cd62L

#### Figure S2

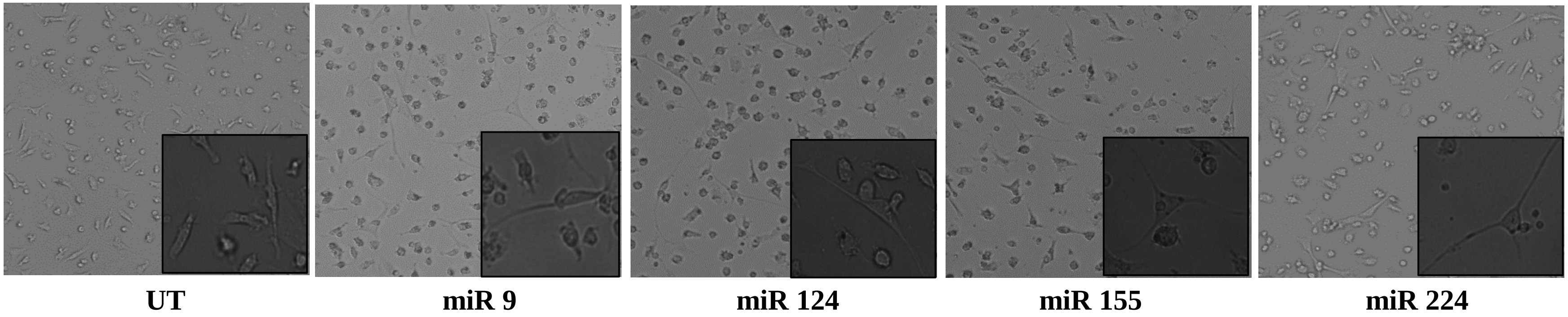


Mouse Bone marrow derived cells were converted to macrophages with M-CSF for 7 days. Cells were transfected with respective miRs for 24 hrs and incubated in NIM for 48 hrs. This protocol was repeated twice and images were captured in Olympus 200X and 400X magnification (insert image). miR transfected cells show high branching and elongated dendritic processes.

#### Figure S3 A

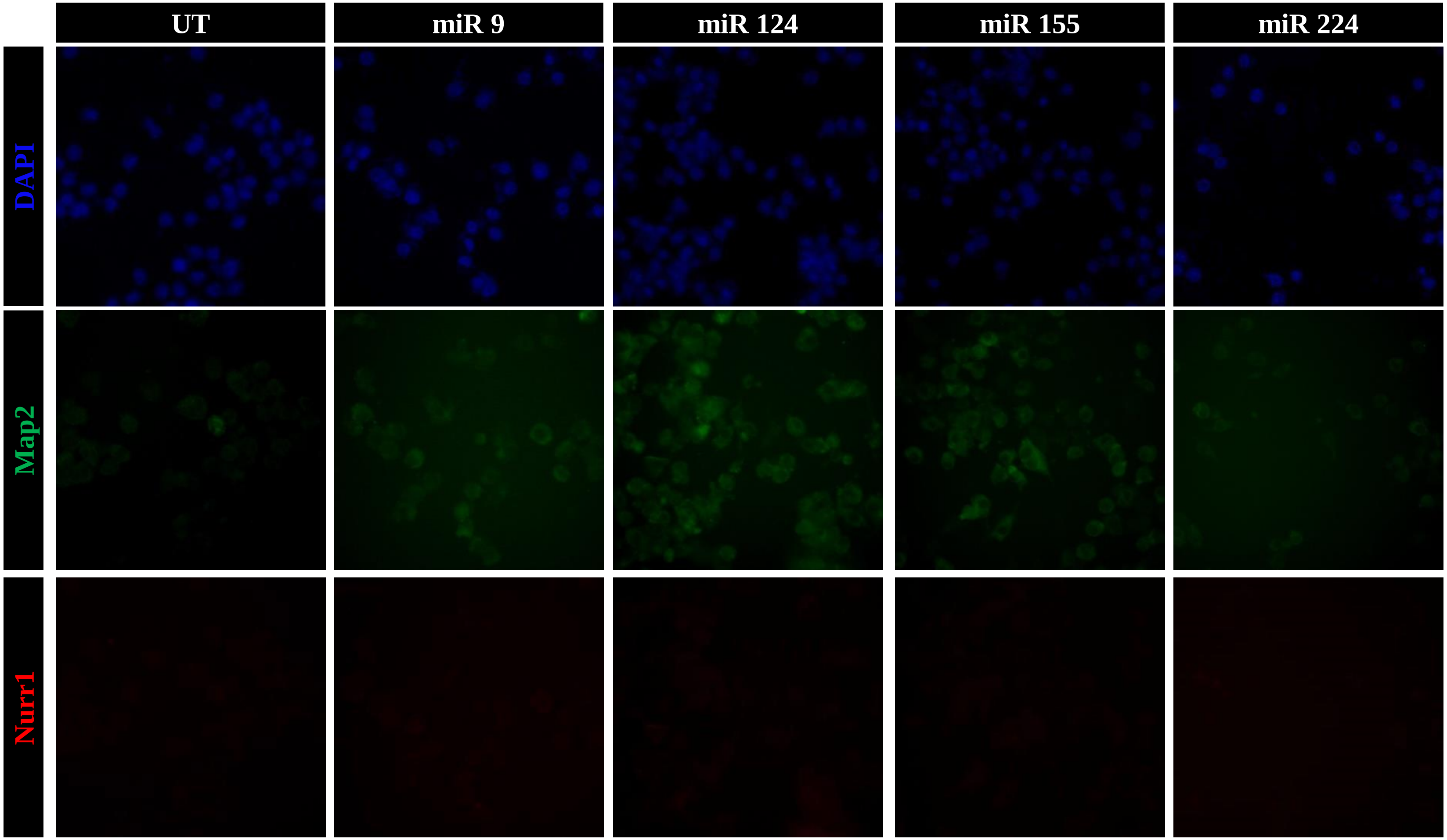


Raw 264.7 cells were transfected with respective miRs thrice for 24 hrs and incubated in NIM for 48 hrs. (A) Cells were stained with DAPI, Map2 and Nurr1; and images were captured in Olympus at 400X magnification.

#### Figure S3

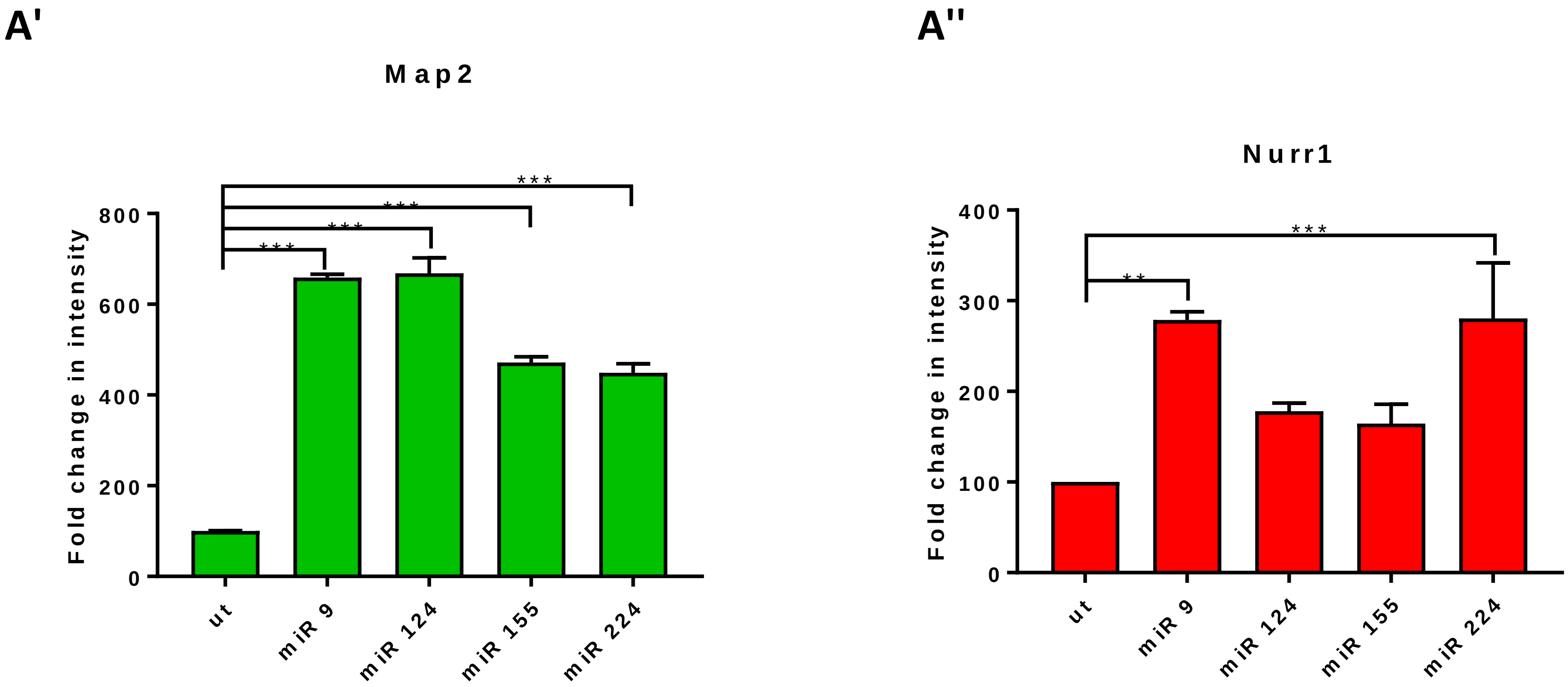


Three individual Immuno-flouresence images of Map2 and Nurr1 stained were processed in Image J for mean intensity. A‘) Map2-FITC fluorescence intensity was plotted as fold change. A‘) Nurr1-Alexa 546 fluorescence intensity was plotted as fold change. *** represents P<0.0001; ** represents P<0.005

**Figure S3 B**

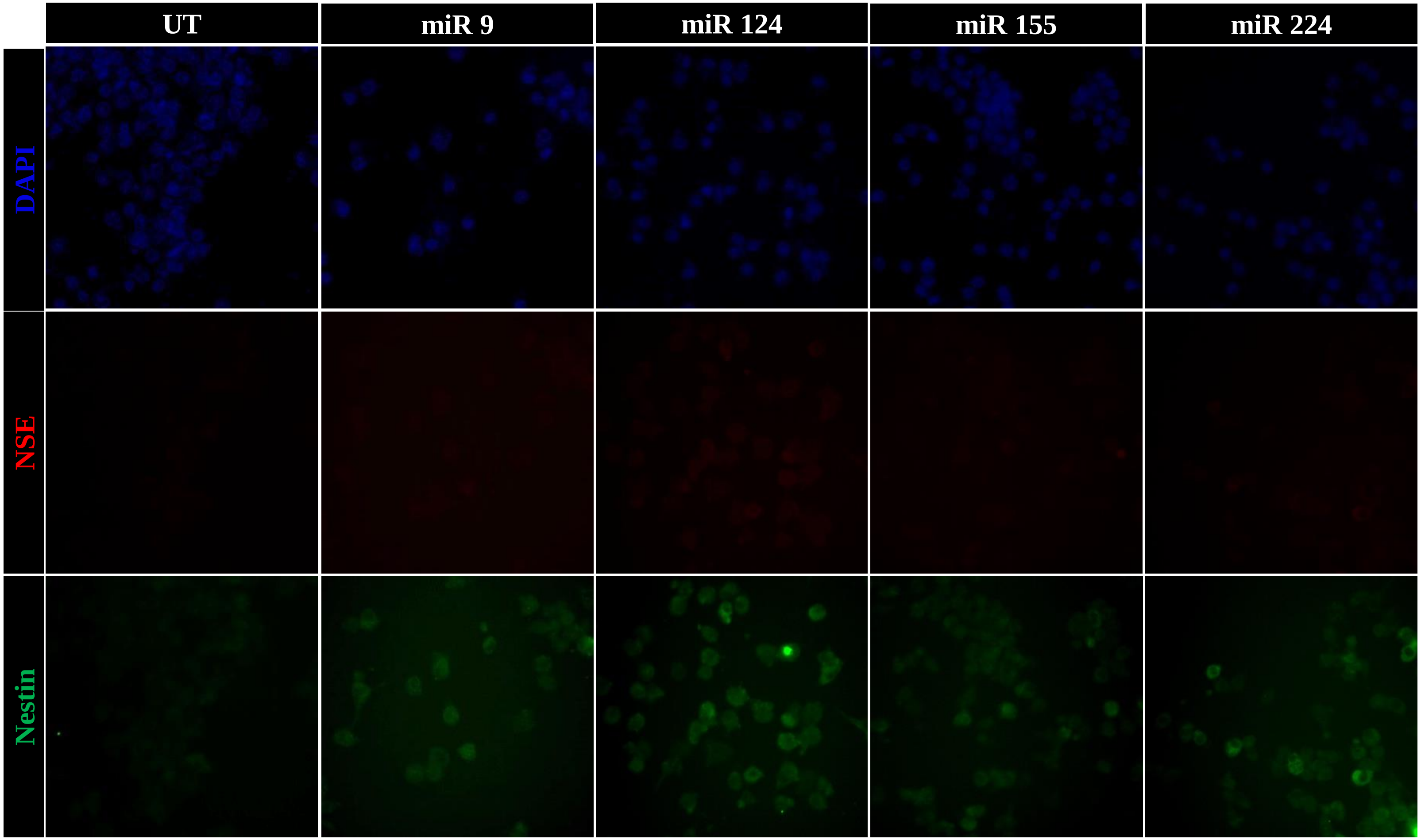


(B) Cells were stained with DAPI, NSE and Nestin; and images were captured in Olympus at 400X magnification.

### Figure S3

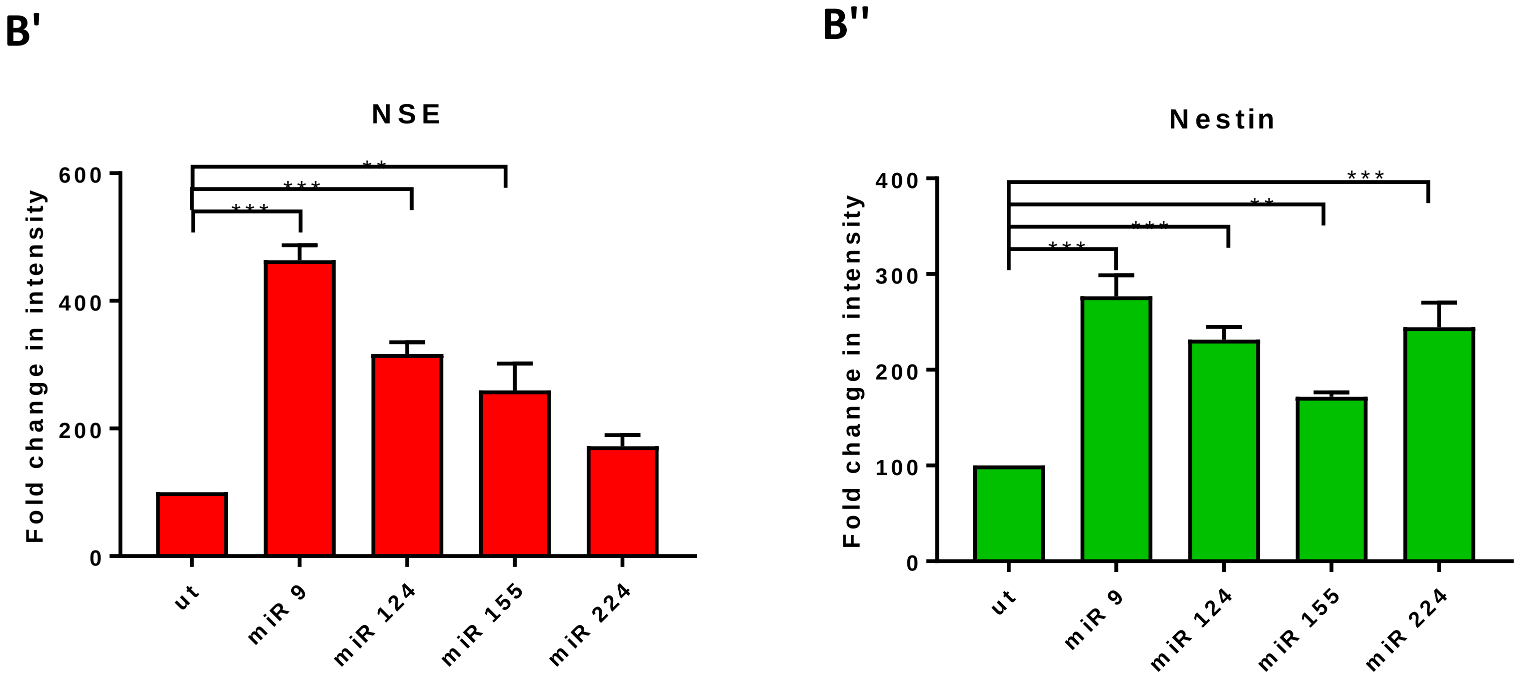


Three individual Immuno-flouresence images of Map2 and Nurr1 stained were processed in Image J for mean intensity. B‘) NSE-Alexa 546 fluorescence intensity was plotted as fold change. B‘) Nstin-Alexa 488 fluorescence intensity was plotted as fold change. *** represents P<0.0001; ** represents P<0.005

**Figure S3 C**

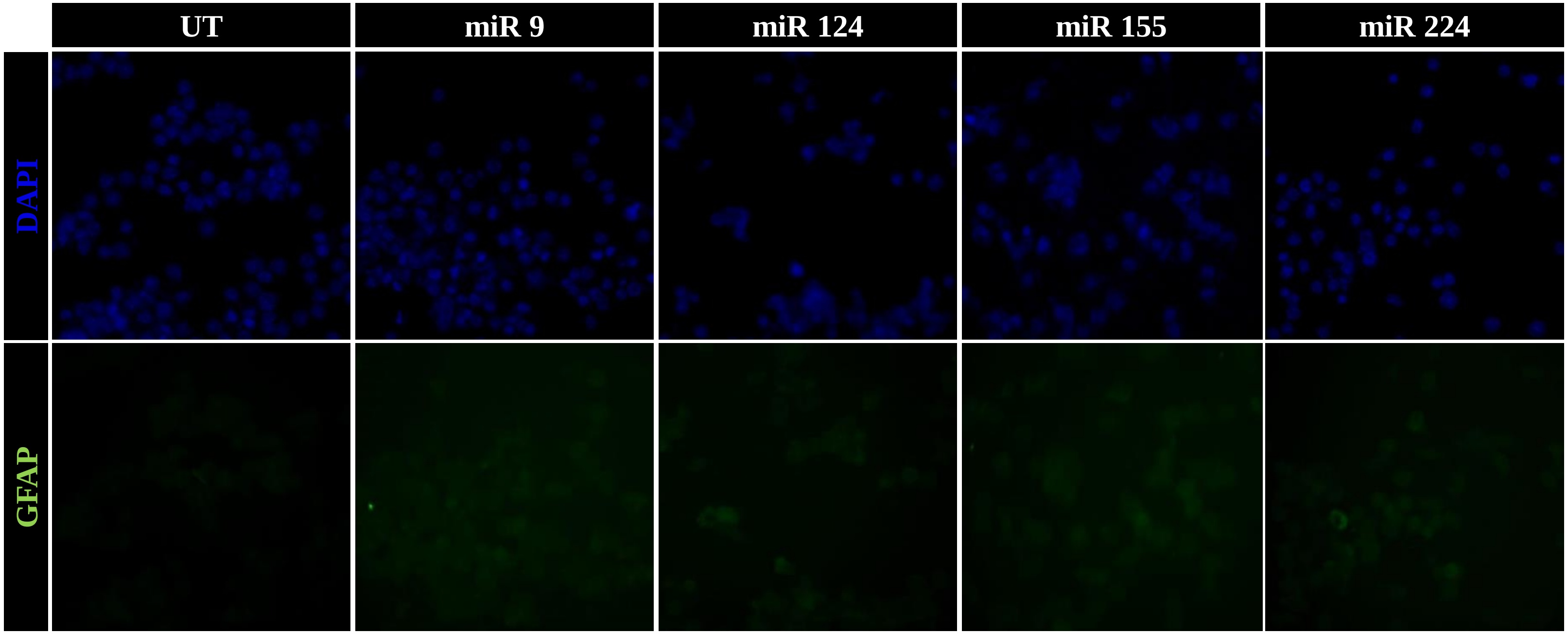


(A) Cells were stained with DAPI and GFAP; and images were captured in Olympus at 400X magnification. .

**Figure S3 D**

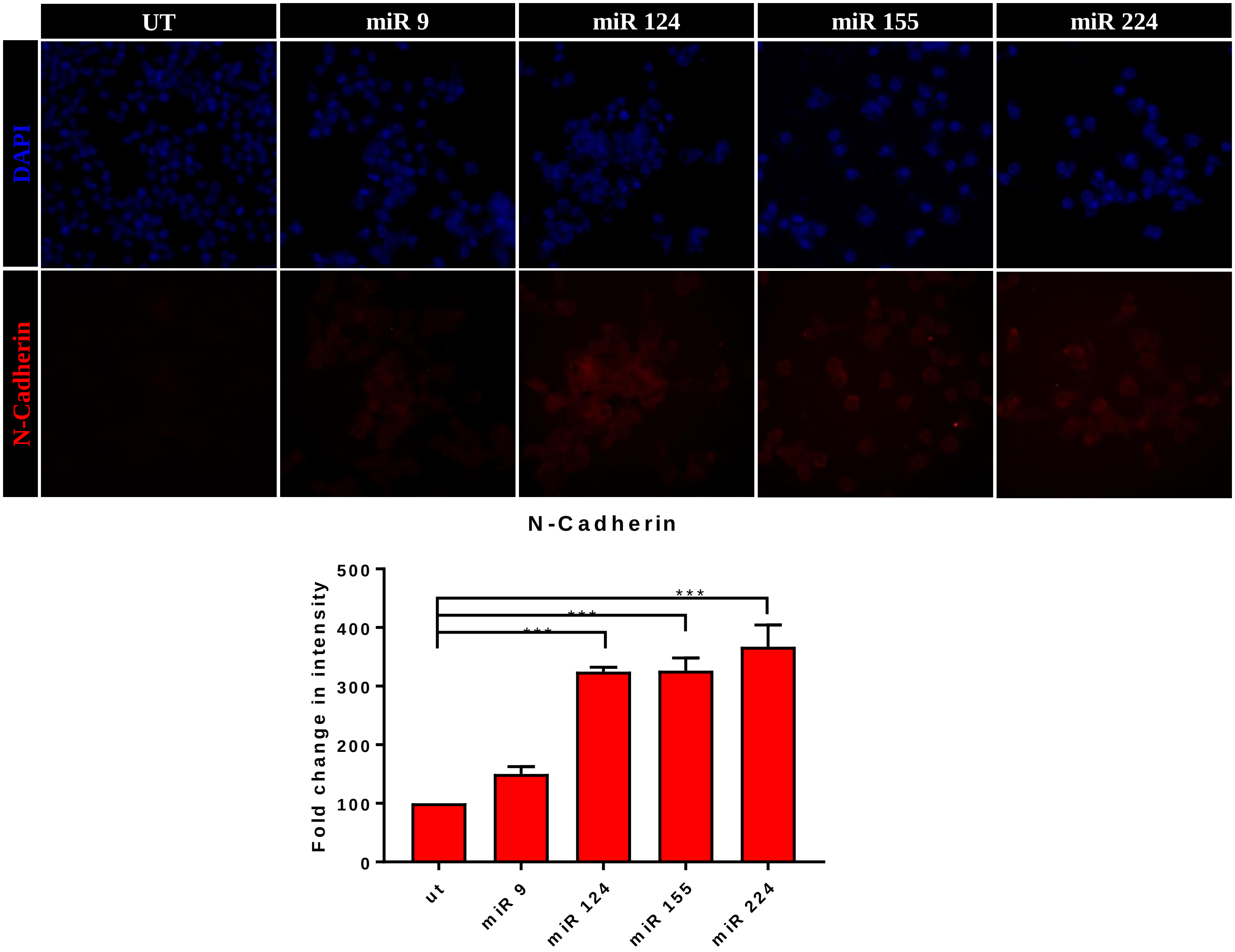


(D) Cells were stained with DAPI and N-Cadherin; and images were captured in Olympus at 400X magnification. D') N-Cadherin-Alexa 546 fluorescence intensity was plotted as fold change. *** represents $P<0.0001$

#### Figure S4

A

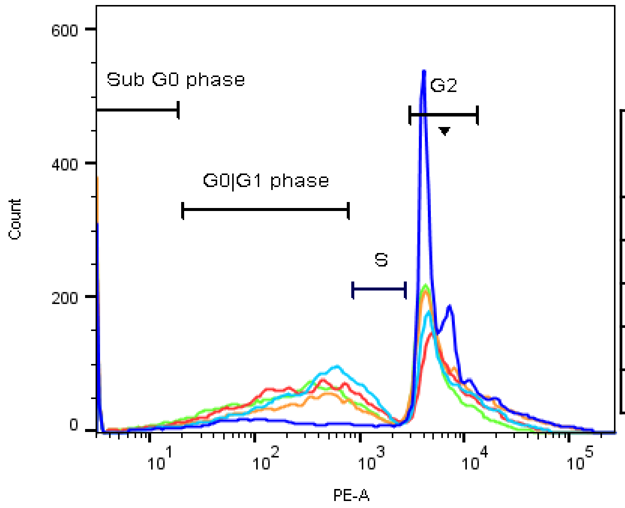


B

| Group | Cell Cycle analysis | | | |
|---|---|---|---|---|
| | Sub G0 (%) | G0/G1 (%) | S (%) | G1/S (%) |
| Un treated | 6.77 | 13.0 | 2.93 | 62.3 |
| miR 9 | 6.11 | 39.6 | 11.7 | 32.6 |
| miR 124 | 7.71 | 42.9 | 9.56 | 30.4 |
| miR 155 | 8.41 | 29.5 | 6.42 | 40.1 |
| miR 224 | 9.15 | 41.3 | 7.06 | 35.6 |

Macrophages were reprogrammed to iNCs and analysed for cell cycle stages through flow cytometry. (A) Different stages of cell cycle of various miR-induced iNCs. (B) % of iNCs in various stages.

**Figure S5**

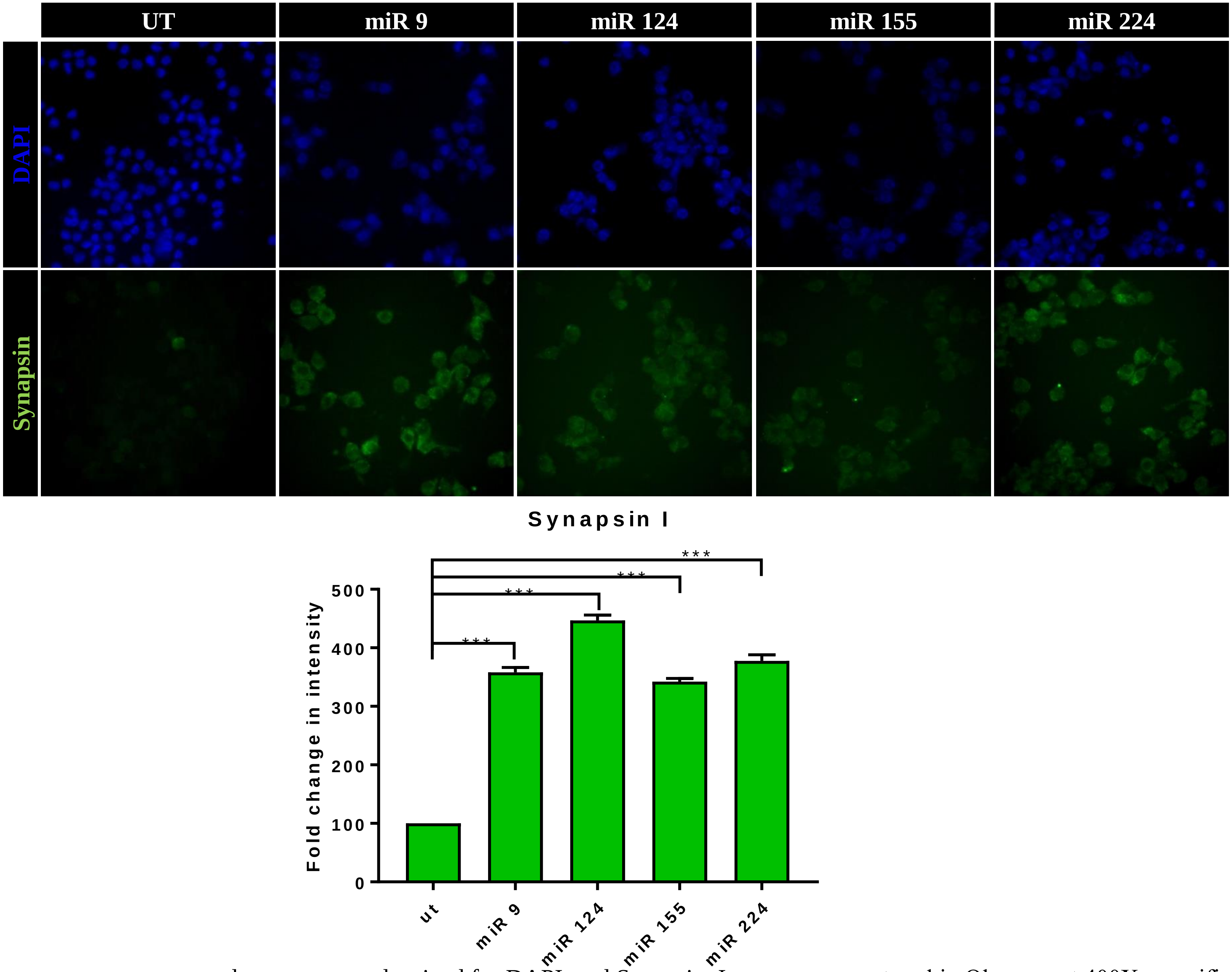


Macrophages were reprogrammed to neurons and stained for DAPI and Synapsin. Images were captured in Olympus at 400X magnification. D‘) Synapsin I-Alexa 488 fluorescence intensity was plotted as fold change. *** represents P<0.0001

### Figure S6

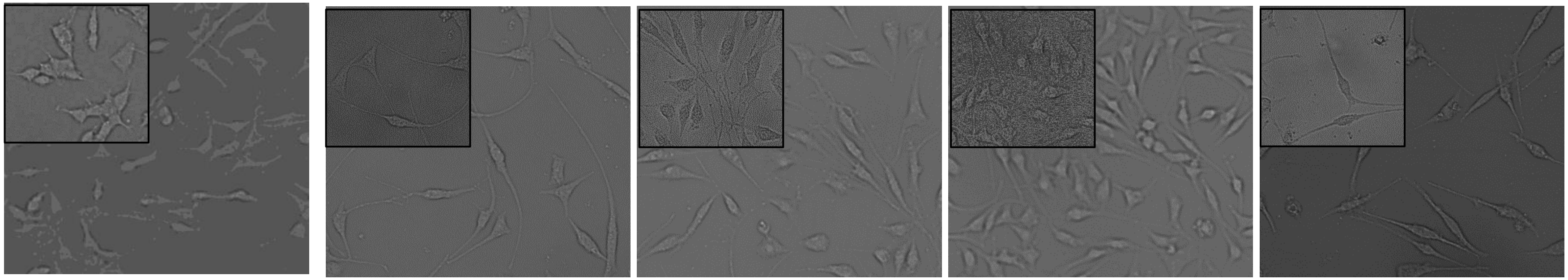

UT miR 9 miR 124 miR 155 miR 224

Mouse Fibrobalst Cell line, L929 were transfected with respective miRs for 24 hrs and incubated in NIM for 48 hrs. This protocol was repeated twice and images were captured in Olympus 200X and 400X magnification (insert image). miR transfected cells show high branching and elongated dendritic processes.

### Supplementary Videos

**SV1:** miR 124 mediated reprogrammed neural cells treated with Fluro 4 AM as mentioned in the protocols. Images were captured during/ immediately after addition of stimulant in FITC filter in florescence microscope for every sec for 30Sec and merged together to form a motion video.

**SV2:** miR 155 mediated reprogrammed neural cells treated with Fluro 4 AM as mentioned in the protocols. Images were captured during/ immediately after addition of stimulant in FITC filter in florescence microscope for every sec for 30Sec and merged together to form a motion video.
