## Supplemental Table for "microRNAs (miR 9, 124, 155 and 224) transdifferentiate macrophages to neurons"

**Supplementary Tables**

| **miRNA** | **Sequence (5'-3')** |
|---|---|
| mmu-miRNA 9-3p | UCUUUGGUUAUCUAGCUGUAUGA |
| mmu-miRNA 124-3p | UAAGGCACGCGGUGAAUGCC |
| mmu-miRNA 155-3p | CUCCUACCUGUUAGCAUUAAC |
| Mmu-miRNA 224-3p | AAAUGGUGCCCUAGUGACUACA |

Table S1: miRNA sequences used in the study

| **Gene** | | **Sequence** |
|---|---|---|
| *miRNA 9* | Forward | ATCTAGCTGTATGAGTGCCACA |
| | Reverse | TCTAGCTTTATGACGGCTCTGT |
| *miRNA 124* | Forward | GCCTCTCTCTCCGTGTTCACA |
| | Reverse | CGCGTGCCTTAATTGTATGGA |
| *miRNA 155* | Forward | ATGCTAATCGTGATAGGGGTTTTT |
| | Reverse | AGGAGTCAGTTGGAGGCAAA |
| *miRNA 224* | Forward | TCAAGTCACTAGTGGTTCCGT |
| | Reverse | ACTAGGGCACCATTTTGAAACA |
| *Sirt 1* | Forward | GAGCTGGGGTTTCTGTTCTCC |
| | Reverse | AACATGGCTTGAGGGTCTGG |
| *Ptx 3* | Forward | GCTCATGCTTTGGAGCGTT |
| | Reverse | GAGGCGAAATTTTGCTTCGCA |
| *LamC 1* | Forward | AGCCGACTGCAGAATATCCG |
| | Reverse | CCCTGGAGGCGATCTCAATC |
| *FADD* | Forward | TGCTCCATCTGGCTGTTTGT |
| | Reverse | AGTCTGGGGAGTCAAGAGCA |
| *MAP2* | Forward | CTCCTCCAAAGTCCCCAGCTA |
| | Reverse | TCCGGCAGTGGTTGGTTAATAA |

| | | |
|---|---|---|
| *Tubb3* | Forward | TCACGCAGCAGATGTTCGAT |
| | Reverse | GTGGCGCGGGTCACA |
| *Mash1* | Forward | GGCAGCAGATCCTGCATCTT |
| | Reverse | TTTTCTGCCTCCCATTTGA |
| *GAPDH* | Forward | CATGGCCTTCCGTGTTCCTA |
| | Reverse | GCGGCACGTCAGATCCA |

Table S2: Primers used in the study for real time PCR

| **Gene** | | **Sequence** | **Annealing Temp.** |
|---|---|---|---|
| *Nestin* | Forward | AAGTTCCCAGGCTTCTCTTG | 49 |
| | Reverse | GTCTCAAGGGTATTAGGCAAGG | |
| *CD11b* | Forward | CATCCCATGACCTTCCAAGAG | 50˚C |
| | Reverse | GTGCTGTAGTCACACTGGTAG | |
| *Pu.1* | Forward | AGAGCTATACCAACGTCCAATG | 50 |
| | Reverse | GTGAAGTGGTTCTCAGGGAAG | |
| *C-Myc* | Forward | GTTGGAAACCCCGCAGACAG | 55 |
| | Reverse | ATAGGGCTGTACGGACTCGT | |
| *NeuN* | Forward | GTAGAGGGACGGAAAATTGAGG | 50 |
| | Reverse | GGGAAACTGGTCACTGCATAG | |
| *NGFR* | Forward | AGAGTATGTCCGCTCCCTGT | 56 |
| | Reverse | CCAGGGATCTCCTCGCATTC | |
| *Sox2* | Forward | CCCCGGGCTGCAGGAAT | 54 |
| | Reverse | CCGGGCTGTTCTTCTGGTT | |
| *Oct 4* | Forward | GCAGATAGGAACTTGCTGGGT | 55 |
| | Reverse | CACCTTTCCAAAGAGAACGCC | |
| *Klf4* | Forward | GCGAGTCTGACATGGCTGT | 55 |
| | Reverse | GTTCCTCACGCCAACGGTTA | |

| | | | |
|---|---|---|---|
| *Nanog* | Forward | AGGCCTGGACCGCTCAGT | 55 |
| | Reverse | AGTTATGGAGCGGAGCAGCAT | |
| *β-Actin* | Forward | GGCTGTATTCCCCTCCATCG | 53 |
| | Reverse | CCAGTTGGTAACAATGCCATGT | |

Table S3: Primers used in the study for sqRT-PCR

| miRNA | miR 9 | miR 124 | miR 155 | miR 224 |
|---|---|---|---|---|
| Unique predicted gene targets pooled from targetscan.7, mirGator, miRecords | *ONECUT2*<br>*ONECUT3*<br>*ONECUT1*<br>*C1orf220*<br>*KIAA0825*<br>*POU2F1*<br>*SLC2A2*<br>*GREB1*<br>*AC003102.1*<br>*CYP3A5*<br>*IFT88*<br>*PPCS*<br>*CHST9*<br>*POU6F2*<br>*EHD4*<br>*PRDM6*<br>*LDLRAP1*<br>*CAMK1D*<br>*RAB34*<br>*C9orf89*<br>*IL1F10*<br>*MDGA2*<br>*TSLP*<br>*ERG*<br>*KCNJ2*<br>*CDR1as*<br>*AP1S2*<br>*LIN28B*<br>*TNC*<br>*RNF146*<br>*C1ORF220*<br>*DNAJC14*<br>*SLC31A2*<br>*LEPRE1*<br>*MDGA2*<br>*C9orf170*<br>*EXPH5*<br>*STK3*<br>*PRRX1*<br>*PBOV1*<br>*TESK2*<br>*TMEM126B* | *CTDSP1*<br>*LRRC58*<br>*SEPT10*<br>*B4GALT1*<br>*TRIB3*<br>*VAMP3*<br>*SLC10A7*<br>*SLC31A2*<br>*RHOG*<br>*RNPEPL1*<br>*HAUS4*<br>*SERINC2*<br>*NOX4*<br>*CEBPA*<br>*SNAI2*<br>*AL626787.1*<br>*SLC50A1*<br>*GCDH*<br>*QKI*<br>*PDCD6*<br>*DNASE2*<br>*KIAA0247*<br>*CHIC1*<br>*SLITRK6*<br>*CYYR1*<br>*APLN*<br>*RAB34*<br>*FLOT2*<br>*PTPN12*<br>*VPS37C*<br>*ASPA*<br>*FAM150A*<br>*CXorf64*<br>*PABPC4L*<br>*PIK3C2A*<br>*TARBP1*<br>*XRCC6*<br>*FOXQ1*<br>*CTXN1*<br>*FOLR1*<br>*RFX4*<br>*CBX2* | *ZNF345*<br>*ANKRD20A2*<br>*C5orf28*<br>*PMAIP1*<br>*GRIA3*<br>*FAM159B*<br>*TSHZ3*<br>*ANKRD20A4*<br>*FOXR2*<br>*EFCAB10*<br>*WWC1*<br>*RP11-382J12.1*<br>*ITGBL1*<br>*STARD6*<br>*PDE6H*<br>*AC107021.1*<br>*SLCO1A2*<br>*RNPC3*<br>*ACTL7A*<br>*ACTA1*<br>*ARID2*<br>*SSBP3-AS1*<br>*C11orf53*<br>*H3F3A*<br>*TRAPPC3L*<br>*RNF123*<br>*UAP1*<br>*DYNC1I1*<br>*PBK*<br>*AADAT*<br>*DET1*<br>*ATXN7L3B*<br>*HNRNPA3*<br>*CEBPB*<br>*HIST2H4B*<br>*ASS1*<br>*NFE2L2*<br>*RPS6KB1*<br>*PICALM*<br>*RAB6A*<br>*PEX5L*<br>*MOCOS* | *KRTAP6-3, SLMO2,*<br>*RABGGTB, PTX3,*<br>*GNG11, CCKBR*<br>*ISM1,*<br>*DDC,*<br>*HEBP1,*<br>*RP11-849H4.2*<br>*ATXN7L3B*<br>*CLEC6A*<br>*FAM27E2*<br>*SMIM14*<br>*FAM27E3*<br>*TMEM133*<br>*FAM27E1*<br>*PCTP*<br>*HOXD10*<br>*SLC13A1*<br>*ZDHHC20*<br>*SPICE1*<br>*SPPL3*<br>*RGS21*<br>*DYM*<br>*TSLP*<br>*WDR55*<br>*C11orf34*<br>*MMGT1*<br>*ZNF174*<br>*DDIT3*<br>*FSCN3*<br>*DENND5B*<br>*LRIF1*<br>*TMEM196*<br>*HSCB*<br>*NAPB*<br>*MICALL1*<br>*RMDN3*<br>*PROSC*<br>*ZNHIT6*<br>*GNG13*<br>*LYG2*<br>*UQCRH*<br>*APITD1* |

| | | | | |
|---|---|---|---|---|
| | *ZNF197* | *PGF* | *CCDC130* | *C1D* |
| | *SLC50A1* | *ACADVL* | *AICDA* | *SLC1A6* |
| | *TCF15* | *OAF* | *CD274* | *CCDC43* |
| | *TGFBI* | *S100A4* | *SELT* | *C8orf46* |
| | *HN1L* | *TSPAN15* | *ZMYM6NB* | *SCRN2* |
| | *AC117834.1* | *MPV17* | *TMPRSS11BNL* | *NDRG3* |
| | *MESDC1* | *SERP1* | *ZMYM6* | *DGKA* |
| | *OPALIN* | *CD164* | *TCF4* | *F2R* |
| | *CALB2* | *MAP3K2* | *LRRCC1* | *SPAG1* |
| | *AL603965.1* | *EGFL6* | *JARID2* | *EIF5A2* |
| | *AL591684.1* | *SLC16A1* | *MYB* | *PDGFRA* |
| | *C10orf91* | *OXNAD1* | *MEIS1* | *TRAPPC3* |
| | *EMB* | *CHP1* | *HBP1* | *CAPN10* |
| | *IFNG* | *FRMD4B* | *DNTTIP1* | *FARP2* |
| | *SLC10A3* | *FA2H* | *CYB561D1* | *RUVBL2* |
| | *UCP1* | *WASF2* | *NR1H3* | *CPNE8* |
| | *SRA1* | *CCBL2* | *RUFY2* | *GCM1* |
| | *ANO1* | *NAT8L* | *ANKDD1B* | *H3F3B* |
| | *MAEA* | *SGPP1* | *C1orf101* | *AC023590.1* |
| | *RNF150* | *LGALS3* | *INADL* | *CD53* |
| | *UTRN* | *PTGES* | *C15orf56* | *TMEM9B* |
| | *LECT1* | *DHFR* | *CXorf38* | *HOXA5* |
| | *MAGT1* | *CCDC177* | *ZIC3* | *RBMXL2* |
| | *SCRIB* | *FZD4* | *C7orf71* | *P2RY4* |
| | *FRMD6* | *PCDH8* | *TP53INP1* | *ZNF774* |
| | *NXPE3* | *MYO10* | *GDAP2* | *CHIT1* |
| | *RANBP17* | *BMP6* | *CCDC152* | *CCDC177* |
| | *RFESD* | *TMEM256-* | *RGP1* | *CROT* |
| | *CAPZA1* | *PLSCR3* | *MAK16* | *RP11-67H2.1* |
| | *CCDC43* | *RAD51L3-RFFL* | *CENPW* | *RAB40B* |
| | *SMARCD2* | *SPOPL* | *MYO1D* | *C20orf96* |
| | *CDRT4* | *PPIL6* | *HLA-DRA* | *ATF6B* |
| | *IPO4* | *TOR3A* | *FGF7* | *RNF5* |
| | *SHROOM4* | *EYA4* | *GDF6* | *ELOVL6* |
| | *DYRK1B* | *PTBP1* | *MEP1A* | *ZBTB14* |
| | *SAMD11* | *CBL* | *SMR3B* | *MAFG* |
| | *TRPM7* | *ZKSCAN3* | *BPIFB2* | *GALR1* |
| | *YPEL2* | *CNEP1R1* | *AARD* | *ASB8* |
| | *PAK4* | *IMP3* | *ZNF652* | *IL12B* |
| | *ATOH8* | *KIAA1671* | *C1QL4* | *C6orf201* |
| | *FAM19A5* | *MARVELD1* | *DHX40* | *OSBPL2* |
| | *FOXP4* | *VAT1* | *RP11-661C8.3* | *ELP6* |
| | *CXCL11* | *SIX4* | *MFSD2A* | *ARF6* |
| | *FBN1* | *PTBP2* | *TRAPPC2* | *CDS2* |
| | *UBE3C* | *GALNT10* | *GRIA4* | *ADH1B* |
| | *ORAOV1* | *KRT3* | *AC023590.1* | *GMPS* |
| | *LGR6* | *RFFL* | *C3orf18* | *LYPD6* |
| | *RHOA* | *STT3A* | *ARL4A* | *PARD3* |
| | *IFRG15* | *NME4* | *MT-ATP8* | *SH2D1A* |
| | *ZNF74* | *MDK* | *MEST* | *KRTAP10-1* |
| | *CCNE2* | *AHR* | *CSF1R* | *LATS2* |
| | *NHSL1* | *ASCC2* | *ZNF488* | *DTNBP1* |
| | *TNP1* | *TMEM104* | *NDN* | *NAP1L5* |
| | *LETM2* | *TUB* | *MGP* | *CDCA5* |
| | *POU2F2* | *C10orf12* | *YIPF7* | *ABHD3* |

| | | | |
|---|---|---|---|
| | *SH3TC2*<br>*SHC2*<br>*NTNG1*<br>*Foxg1* | *RARG*<br>*SNTB2*<br>*PGRMC2*<br>*PNPLA2*<br>*LOC*<br>*ITGB1*<br>*Sycp1*<br>*Foxa2*<br>*Mtpn*<br>*LMNB1*<br>*c14orf24*<br>*JAG1*<br>*DLX2*<br>*SOX9*<br>*MCT1*<br>*Lhx2* | *E2F2*<br>*INPP5D*<br>*ARMCX2*<br>*Spfi 1*<br>*Rheb*<br>*bat5*<br>*FADD*<br>*IKBKE*<br>*Ripk1*<br>*socs1*<br>*kgf*<br>*FOS*<br>*HiF1a*<br>*cd62L*<br>*LOC424442*<br>*ITGB1*<br>*Ctdsp1*<br>*Ptbp1*<br>*Ptbp2*<br>*Sycp1*<br>*Foxa2*<br>*Mtpn*<br>*Itgb1*<br>*LAMC1*<br>*PTPN12*<br>*LMNB1*<br>*c14orf24*<br>*JAG1*<br>*DLX2*<br>*SOX9*<br>*MCT1*<br>*Lhx2* |
| Common genes | *SLC31A2, SLC50A1,* | | *ATXN7L3B* |
| | | *Ctdsp1* | |

Table S4: miRNA predicted targets. Top 100 genes from miRGator, TargestScan, miR records, were pooled and added in each row. Genes manipulated by each miR target alone were grouped in a unique predicted row. Genes manipulated by more than one miR are added in common genes row.

Equitation used to calculate efficacy of reprogramming from FACS data.

$$Efficay\ of\ reprogramming\ (\%) = \left\{ \frac{\left( \frac{Cells\ obtained\ after\ 3\ transfections}{Total\ no.of\ macrophages\ seeded} \right) \times 100}{100} \right\} \times purity\ of\ neurons(Q3)$$
